# A Toolbox for Biomanufacturing of Functionalised PHA Nanoparticles with *C. necator*

**DOI:** 10.64898/2026.03.17.712365

**Authors:** John Allan, Lisa J. K. Zillig, Simona Della Valle, Harrison Steel

## Abstract

Microbes have the potential to manufacture plastics from sustainable feedstocks while enabling novel material properties and functions that are not easily accessible through conventional chemical synthesis. Realising this potential requires a comprehensive genetic and process engineering framework that spans chassis and bioprocess optimisation, polymer property control, and downstream functionalisation. Here we develop such a platform in *Cupriavidus necator,* with a focus on high-value polyhydroxyalkanoate (PHA) nanoparticles. To this end we first optimise the transformation protocol for the organism. Next, we create a library of PhaC synthase variants from *C. necator*, *Aeromonas caviae* and *Brevundimonas sp.* in a ΔphaC background, demonstrating that they allow customisation of the material properties of produced PHA particles. Our results combine data from Flow cytometry, Transmission Electron Microscopy (TEM), Fourier Transform InfraRed Spectroscopy (FTIR), and Differential Scanning Calorimetry (DSC) to show that it is possible to generate materials ranging from highly crystalline PHAs to softer P(3HB-co-3HHx) copolymers and that an *A. caviae* PhaC variant can double the yield of large PHA granules. To improve bioprocess sustainability, we coupled *C. necator* with *B. subtilis* in sucrose-fed co-cultures, using tetracycline tolerance differences and inoculation ratios to enhance PHA production from inexpensive, sugar-rich feedstocks. Finally, we add function to the produced PHA nanoparticles by using the molecular protein-fusion technology SpyTag-SpyCatcher, showing it is possible to efficiently capture SpyCatcher-GFP on PHA granules as a proof of concept for PHA’s use as a customisable bio-based nanoparticle. Together, our work offers an innovation to produce bio-PHA nanoparticles in a customisable way, with potential applications in sustainable biomanufacturing, biosensing, drug delivery and future bioremediation technologies.

## Introduction

Producing plastic products sustainably via a biological route without burdening our environment has been proposed as one solution to our global plastic problem (*1–4*). For example, bioplastics can be produced from nature-based resources and are often also biodegradable. One such example is the family of Polyhydroxyalkanoates (PHAs), a polyhydroxyester synthesised by bacteria and plants (*5*, *6*). PHAs have thermoplastic properties, making them amenable to injection moulding and potential replacements for petrochemical plastics like polypropylene (*7*). They are further attractive because they are biocompatible (*8*) and have been explored as materials in various medical implants such as sutures (*9*) or bone grafts (*10*).

Bioplastics can also provide a more sustainable and cost-efficient route to produce high value products like functionalised Nanoparticles (NPs). NPs created with a wide range of compounds (often high-cost materials like gold) are used in diverse applications ranging from medical to agriculture (*11–13*). Creating functional nanoparticles requires a diversity of engineering solutions, spanning their manufacturing over functionalisation to delivery. In this work we focus on developing PHA molecules as a novel chassis for custom-designed environmentally friendly NPs.

To achieve this, we leverage *Cupriavidus necator* H16, an emerging workhorse organism for industrial bioprocessing. It is a Gram-negative mesophile with soil or freshwater habitats that possesses promising metabolic properties: The ability to grow on CO_2_ (*14*) or other gases and organic acids make it an intriguing platform for metabolic engineering, giving the potential to turn waste products into value compounds (*15*).

Combined with the potential of being genetically engineered, *C. necator* presents as an exciting platform organism for biomanufacturing, as demonstrated by its utilization in the production of biofuels (*16*), plant growth promoting hormones (*17*) or high value materials like terpenoids and alkenes (*18*). Further, *C. necator* has already been proven to be able to produce PHA from carbon-rich waste streams such as raw and partially digested agricultural wastes (*19–21*). PHA is a common carbon and energy-storage in bacteria. One of the most dominant forms of PHA is poly-3-hydroxybutyrate (PHB). PHA is synthesised from acetyl Co-A in a three-step metabolic reaction involving the enzymes PhaA, PhaB and PhaC. PhaA forms acetoacetyl-CoA from two molecules of acetyl CoA. PhaB reduces acetoaetyl-CoA to 3-hydroxyacetly-CoA before PhaC polymerises the 3-hydroxyacetyl-CoA monomers into PHA (*22*). The genes, encoding these three enzymes, are organized as a unit in the “phaCAB” operon (*23*). The accumulated PHA polymers are insoluble in water and therefore form granules, with their often large size and complex organisation earning them the status as a distinct organelle, called the *carbonosome* (*24*).

In recent years it has been discovered that various classes and subtypes of PhaC (the enzyme responsible for polymerisation of PHA molecules) exist. The four main classes, I, II, III and IV are distinguished by their different substrate specificities and structural properties (*25*, *26*). While class I, III and IV prefer to incorporate short-chain-length PHA (SCL-PHA) of 3–5 carbon units, class II PhaCs favour medium-chain-length PHA (MCL-PHA) of 6–14 carbon units. On top of this, class I and II are single-unit enzymes, whereas class III and IV both have two different subunits (*27*, *28*). The unique characteristics of each PhaC enzyme impact the properties of the resulting PHA molecule and are therefore a useful target for engineering to determine NP properties (*29*). A versatile and commonly used PHA form is the so-called P(3HB-co-3HHx) copolymer. This represents a mix of the SCL monomer P(3HB) and the MCL monomer 3HHx, in which the long alkyl side chain of 3HHx prevents crystallization. The resulting P(3HB-co-3HHx) becomes more flexible, softer and easier to process for downstream applications with increasing 3HHx composition (*30*).

Various attempts at optimizing the P(3HB-co-3HHx) copolymer composition for bioplastic production have been undertaken. For example, PhaCs with high 3HHx substrate specificity were introduced into recombinant strains (*31*, *32*), and in vitro mutagenesis was harnessed to optimize PhaC enzymes on the background of utilizing a certain carbon resource as starting point for the bioplastic synthesis (*33*). The swift biodegradability in aerobic and anaerobic conditions in comparison to conventional plastics further strengthen the case for P(3HB-co-3HHx) use. On top of this, biocompatibility makes P(3HB-co-3HHx) an ideal material for medical applications (*27*).

Following PHA polymerisation, the PhaC enzyme remains covalently attached to the end of polymer chains (*34*, *35*); this presents an opportunity for other proteins to dynamically interact with PhaC, opening additional possibilities for functionalisation of the system. Past work has demonstrated this by producing fusion proteins which attach enzymes or fluorescent reporters to the surface of purified granules (*34*, *35*), and future development could extend this by (for example) fusing PhaC with a metal-binding or a toxin-degrading enzyme to be deployed in bioremediation.

With higher value than conventional plastic, nanoparticles pose a more attractive product for biomanufacturing. To this end, we have investigated how *C. necator* may be developed as a chassis organism specialised for the purpose of bio-PHA-NP production. This includes the optimisation of the transformation, exploration of producing PHA granules of diverse sizes and polymeric properties to fit diverse applications, sustainable biomanufacturing using co-cultures, and the addition of different functional groups through the engineered SpyTag/SpyCatcher system coupled to PhaC. A key advantage of this modular system is that one well-characterised PHA granule producer strain can be combined with different SpyCatcher-tagged binder-producing partner strains to tune material properties, avoiding repeated genetic re-engineering of the granule producer and enabling mix-and-match tailoring of the bioplastic nanoparticles (Fig. 1).

**Figure 1.**
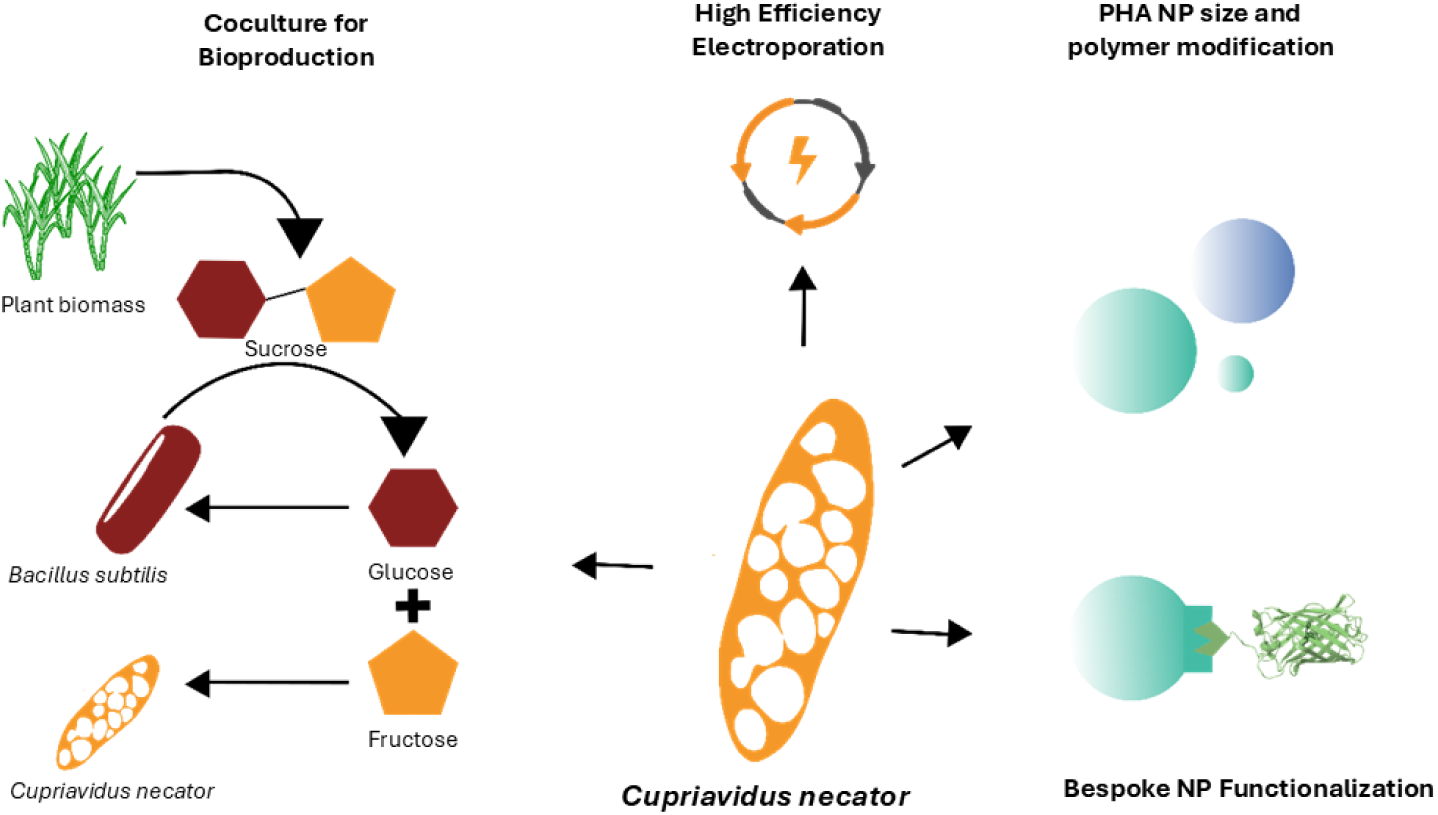
Cupriavidus necator as platform organism for PHA and PHA-nanoparticle production. We present a toolbox for the bacterium Cupriavidus necator, from its transformation to performance in co-culture system, allowing the expansion of substrates for bioproduction and PHA granules modifications into functional nanoparticles for a large and diverse range of applications in medical bioscience and biotechnology.

## Results

### Optimization of Electroporation Transformation of *C. necator*

The first step of our work was to create an efficient and reproducible transformation protocol for *C. necator*. To do this we optimise transformation parameters including OD, outgrowth time, transformation field strength, antibiotic use, media and DNA.

Electrocompetent *C. necator* cells were prepared from pre-cultures spanning an optical density (OD) between 0.2–5 to test whether high-density harvest improves transformation efficiency, expanding on previous work that focused on early to mid-exponential phase (OD 0.4–0.6) cultures (*36*). The baseline electroporation protocol used during optimisation was performed at room temperature using 0.2 cm cuvettes, a field strength of 12.5 kV/cm, 250 ng of the plasmid DNA pSEVA231 (pBBR1, kanR), and a 2 h outgrowth time in super optimal broth (SOB). We first tested the impact of different concentrations of the selection antibiotic kanamycin in the Lysogeny broth (LB)-agar outgrowth plates; concentrations of either 100 or 200 µg/ml kanamycin showed no difference in the formation of colony forming units (CFU) after the transformation (Fig. 2 A). Transformation efficiency increased approximately linearly with culture OD and reached a maximum of (5.8 ± 0.4) × 10^7^ CFU/µg DNA for near-saturated cultures (OD 5), representing an improvement of about two orders of magnitude over earlier reports using related plasmid systems (*36*) (Fig. 2 A), while transformation frequency remained in the range of 10^-3^ - 10^-2^ CFU/cell and was independent of culture density.

**Figure 2.**
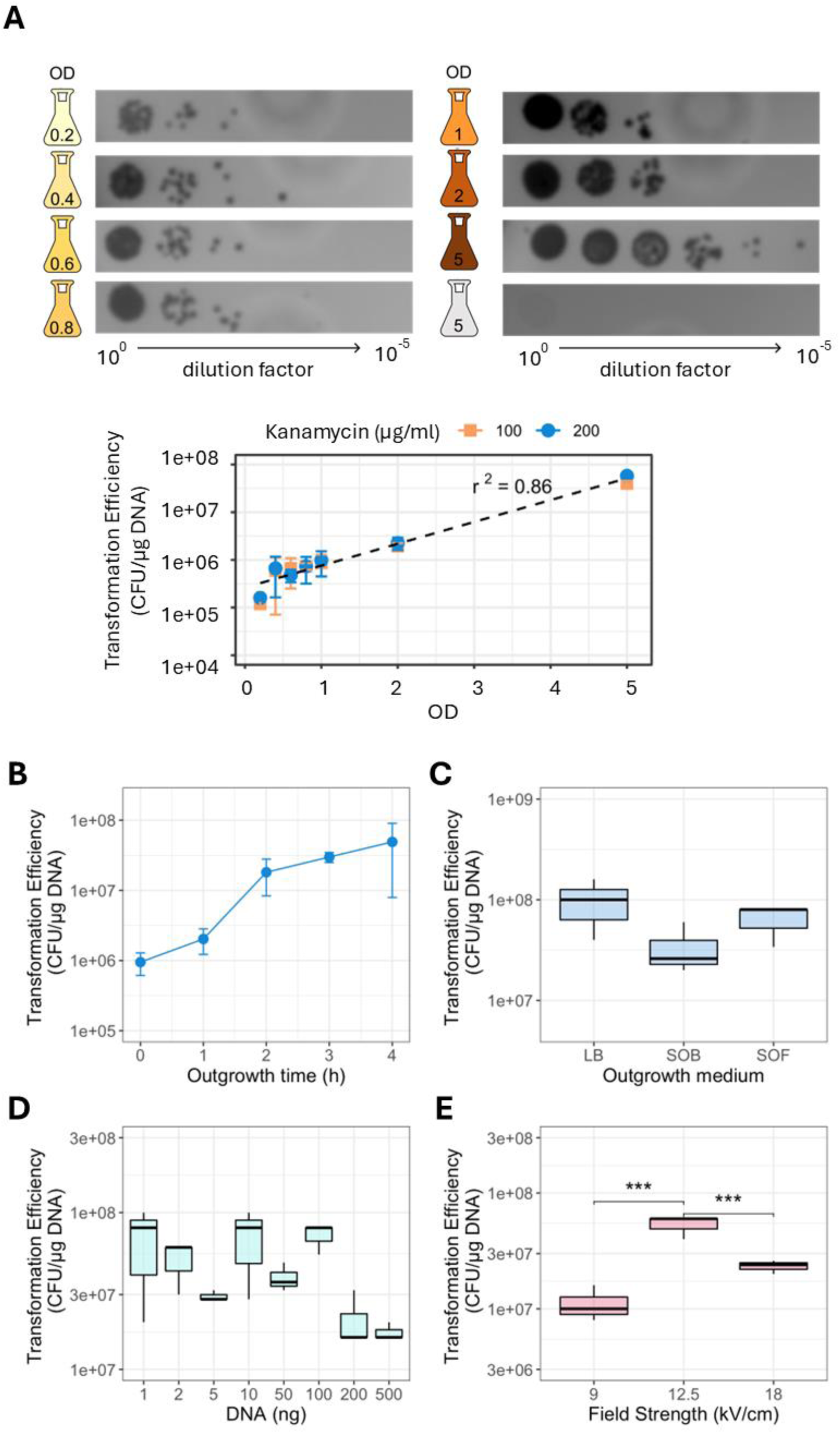
Optimisation of electroporation protocol parameters. **A)** Spot microdilutions of transformation cultures on LB agar solid medium, supplemented with 200μg/ml kanamycin, which were used to quantify transformation efficiency and frequency. For each microdilution, the OD of the cell culture used to prepare the competent cell aliquots is indicated within the flask icon. The last dilution (grey icon) corresponds to a cell aliquot prepared from an OD = 5 culture, where no DNA was added prior to electroporation (i.e., negative control). Transformation efficiency is shown in the graph. **B)** Transformation efficiency as a function of outgrowth duration. Individual points represent the mean ± one standard deviation of n = 3. **C)** Transformation efficiency as a function of recovery medium composition. LB = Luria-Bertani broth, SOB = super optimal broth, SOF = super optimal broth supplemented with 20mM fructose. **D)** Transformation efficiency as a function of DNA mass added to competent cell aliquots**. E)** Transformation efficiency as a function of electroporation field strength. *** = p<0.01, two-sample t-test.

Transformation efficiency increased with outgrowth time up to 2 h but did not improve further at 3–4 h and remained in the order of 10^6^ CFU/µg DNA even without an outgrowth step (Fig 2 B). The choice of recovery medium (LB, SOB, or SOB supplemented with fructose) had no significant effect on transformation efficiency, suggesting that recovery medium composition is not a limiting factor (Fig 2 C). The method was robust across a range of field strengths, all of which outperformed previously reported efficiencies, with 12.5 kV/cm identified as optimal field strength (Fig. 2 E). Finally, transformation efficiency was saturated at very low DNA inputs. No differences were observed between 1–100 ng plasmid DNA, and a reduction in efficiency occurred only at 200–500 ng. This is in direct contrast with other proposed methods and indicates that even low DNA concentrations are sufficient for high electroporation efficiency with the proposed protocol (Fig. 2 D) (*36*, *37*).

Taken together, our results show that an increase in OD in combination with an outgrowth time of 2 h, a field strength of 12.5 kV/cm and moderate DNA (1-100 ng) and antibiotic concentrations (100 µg/ml kanamycin) proved ideal for *C. necator* transformations.

### Generation of PhaC library

Different variants of PhaC from different organisms each possess their own characteristics in catalyzation efficiency and substrate preferences, and past work has modified these attributes further through introduction of mutations (*30*). From this literature we selected a series of variants to screen and expressed these on the plasmids described in Table 1. Different PhaC variants from different organisms, some with, and some without additional mutations were cloned into the pRH412 backbone and electroporated into *C. necator* str H16 ΔphaC (for full protocol see Methods). In total, eight constructs were generated (including the empty vector control) with PhaC variants from three different organisms (*C. necator, A. caviae and B. sp.*). The pRH412 backbone was selected, which encodes an arabinose inducible expression system with a wide range of sensitivities to arabinose and provides reliable and robust gene expression (*38*, *39*).

**Table 1.** PhaC enzyme library for optimization of PHA production. Each PhaC variant has their individual properties leading to changes in overall PHA yield or the fractions of the monomeric units used for polymer production.

| Plasmid name | phaC Origin Organism | Mutations | Properties | Ref. |
| --- | --- | --- | --- | --- |
| pPhaC0 | None – empty vector | na | No PHA production | (38) |
| pPhaC_Cn_wt | *Cupriavidus necator* | none | Wild type phaC | (40) |
| pPhaC_Cn_m | *Cupriavidus necator* | F420S | Increased production | (41) |
| pPhaC_Ac_wt | *Aeromonas caviae* | none | Increased co-polymer production (poly-3HB-co-3HHx) | (42) |
| pPhaC_Ac_m1 | *Aeromonas caviae* | N149S | Increased co-3HHx fraction* | (43–45) |
| pPhaC_Ac_m2 | *Aeromonas caviae* | D171G | Increased co-3HHx fraction | (43–45) |
| pPhaC_Ac_m3 | *Aeromonas caviae* | N149S<br>D171G | Increased co-3HHx fraction | (46) |
| pPhaC_Bs_wt | *Brevundimonas sp.* | none | Trimeric enzyme, functional over wider array of conditions | (47) |

### Quantitative and Qualitative PHA Granule analysis

First, we measured PHA production in *C. necator* with a chromosomal phaC deletion (ΔphaC) expressing these plasmids in the presence and absence of 2 mM arabinose. We used Nile Blue staining in Flowcytometry to quantify the PHA amount. Nile Blue is a lipophilic dye that accumulates in intracellular inclusions like PHA granules and is measurable via fluorescence. Cells stained with this dye were assumed to be PHA positive, and the cells surpassing the control threshold were quantified via FlowJo (Fig. 3 A). The results show that the arabinose reliably induces PHA production at 2mM. Only the *C. necator* H16 wild-type strain without the phaC deletion shows PHA production without arabinose induction, while the negative control with the empty vector (pPhaC0) shows no PHA production, with or without arabinose.

**Figure 3.**
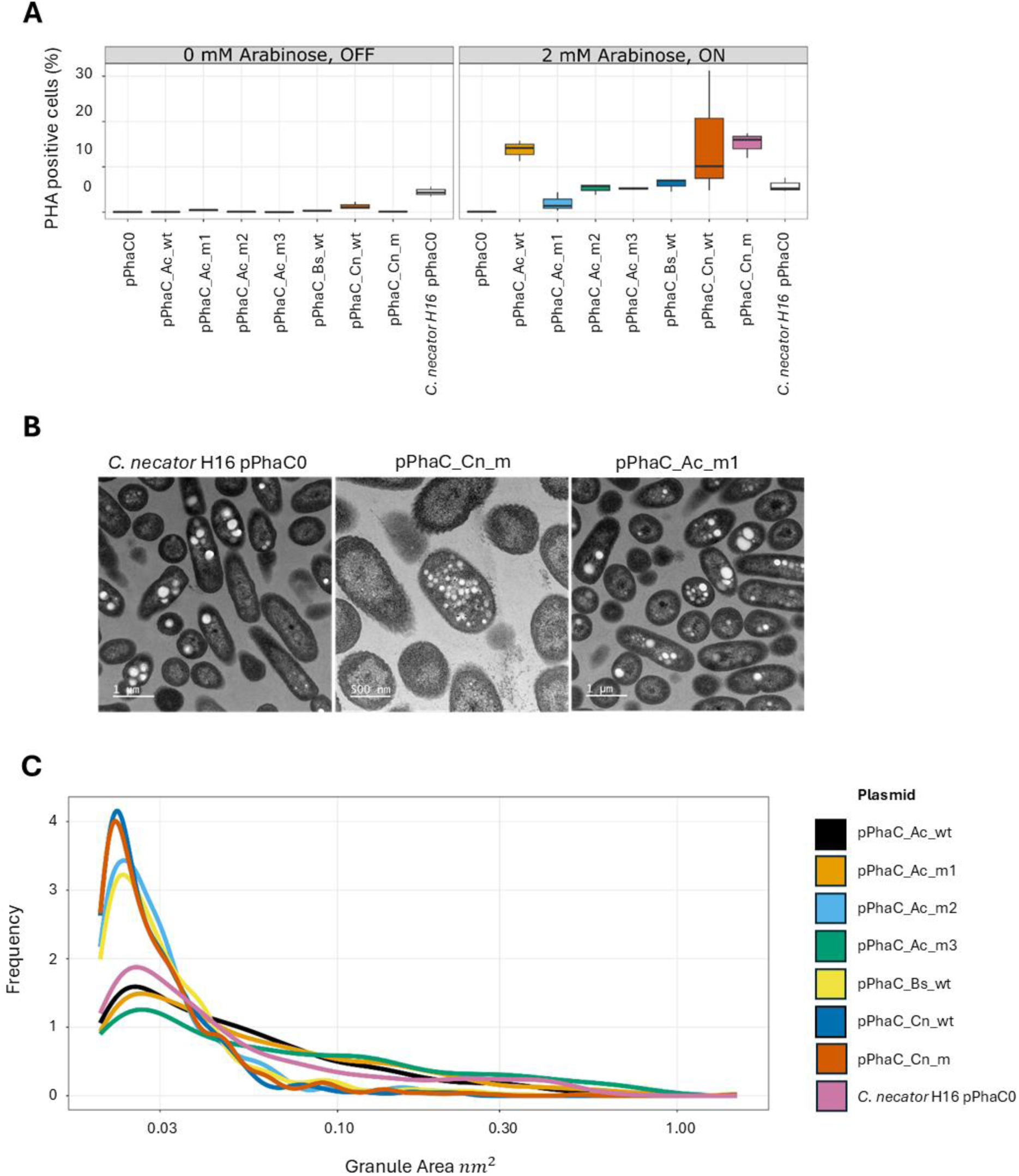
Inducible PHA Production via PhaC Library. **A) FACS:** Validation of C. necator WT empty vector control, which shows stable PHA production regardless of arabinose, as genomic phaC remains. The ΔphaC knockout with empty plasmid produces no PHA. Enzyme variants enable reliable inducible PHA production, with A. caviae, C. necator WT and C. necator mutated variants yielding ∼15% PHA-positive cells. **B) TEM Images left to right**: empty pPhaC0 (few large granules), C. necator mutated pPhaC_Cn_m (small uniform granules, high quantity), A. caviae mutated pPhaC_Ac_m1 (intermediate). Granule amount/size varies by PhaC enzyme. **C) TEM probability distribution of PHA granules.** The Probability Distribution of granule area (nm²) shows most plasmids produce small granules (∼0.02–0.03 nm² peak). Engineered variants (C. necator, A. caviae m2 (D171G mutation), B. sp.) yield sharper peaks and smaller, more uniform granules than WT controls.

The total PHA produced varies between the strains; the most productive enzyme variants were from *A. caviae* or *C. necator* wild type PhaCs, and the *C. necator* mutated variant that has a F420S substitution (pPhaC_Cn_m); each of these variants has approximately 15 % PHA positive cells. Interestingly, the substitution with the plasmidic *C. necator* PhaC (pPhaC_Cn_wt) in the KO strain shows higher PHA production than the WT version, which may be caused by different transcription strength between the genomic and plasmid-borne version (which presents the gene at higher copy number).

Following PHA quantification, the size and frequency of the PHA granules was investigated using Transmission-electron microscopy (TEM) (Fig. 3 B). A supervised machine learning classifier was deployed to analyse the images, identifying cells and PHA granules within them and demonstrating that the size and quantity differ between the variants in the PhaC library. Here, the PhaC enzymes from pPhaC_Cn_wt and pPhaC_Cn_m produced the smallest granules with mean section areas of ∼0.03 nm^2^, and the *A. caviae* double mutant (N149S & D171G; pPhaC_Ac_m3) the largest with section areas as large as ∼0.8 nm^2^ (Fig. 3 C). The cause of the smaller granule size of the pPHaC_Cn_m variant may be a result of the overexpression, where increased quantity of the enzyme in the cell with a similar level of substrate may initiate the production of more, smaller PHA granules compared to the wild type. The PhaCs from *A. caviae* in the wild type version (pPhaC_Ac_wt) and the mutant variants with the N149S mutation (pPhaC_Ac_m1) and the N149S & D171G double mutant (pPhaC_Ac_m3) show higher quantities of the large granules than the WT and especially the other enzyme variants tested (Fig. 3 C).

Next, we investigated the properties of the PHA polymers with Fourier Transform Infrared Spectroscopy (FTIR). This technique uses infrared light to produce a unique spectral fingerprint of a compound, showing the chemical bounds present in the sample. Consequently, the composition of hetero polymers like the P(3HB-co-3HHx) PHA, consisting of different short and medium length chains, can be identified. Previously, *A. caviae* PhaC was observed to produce PHA polymers with enhanced co-3HHx fraction. This polymeric composition is attractive as it increases the elongation break strength (*48*), describing how much stretching a compound can endure before breaking. Past work also showed it was possible to introduce mutations into this PhaC variant that increase the co-3HHx fraction. (*27*, *33*)

For FTIR analysis, *C. necator* cells were cultured in 50 ml batch cultures for 24 h in PHA-M9 (see Methods). Bulk PHA was extracted using the sodium hypochlorite method of Jendrossek (2007) (*49*). The samples were then analysed by FTIR to probe the molecular properties of the polymers. The peaks present in the spectra (Fig. 4) are typical for PHA, arising due to specific functional groups present in the polymer. Peaks around 2934 cm⁻¹ are typically associated with C–H stretching vibrations (methyl and methylene groups) in the polymer backbone. The prominent peak around 1724 cm⁻¹ is the key marker peak for PHAs, corresponding to the carbonyl (C=O) stretching vibration of the ester bond in the polymer. PHA molecules produced by all of the tested strains show a shoulder in the carbonyl peak in comparison to the control sample (LUT) (Fig. 4), indicating a higher fraction of 3HHx in the polymer (*50*). Further peaks at 1458 cm⁻¹ are generally assigned to the asymmetric bending of CH₂ groups, while peaks around 1381 cm⁻¹ correspond to the symmetric bending of terminal CH₃ groups in the polymer; peaks at 1280 cm⁻¹ are assigned to C–O stretching vibrations and related C–O/C–C bounds in the polyester backbone (*51*). The peaks at 1185 cm⁻¹ can be attributed to C–O–C stretching vibrations (*52*, *53*) in the ester linkage in PHA (*54*).

**Figure 4.**
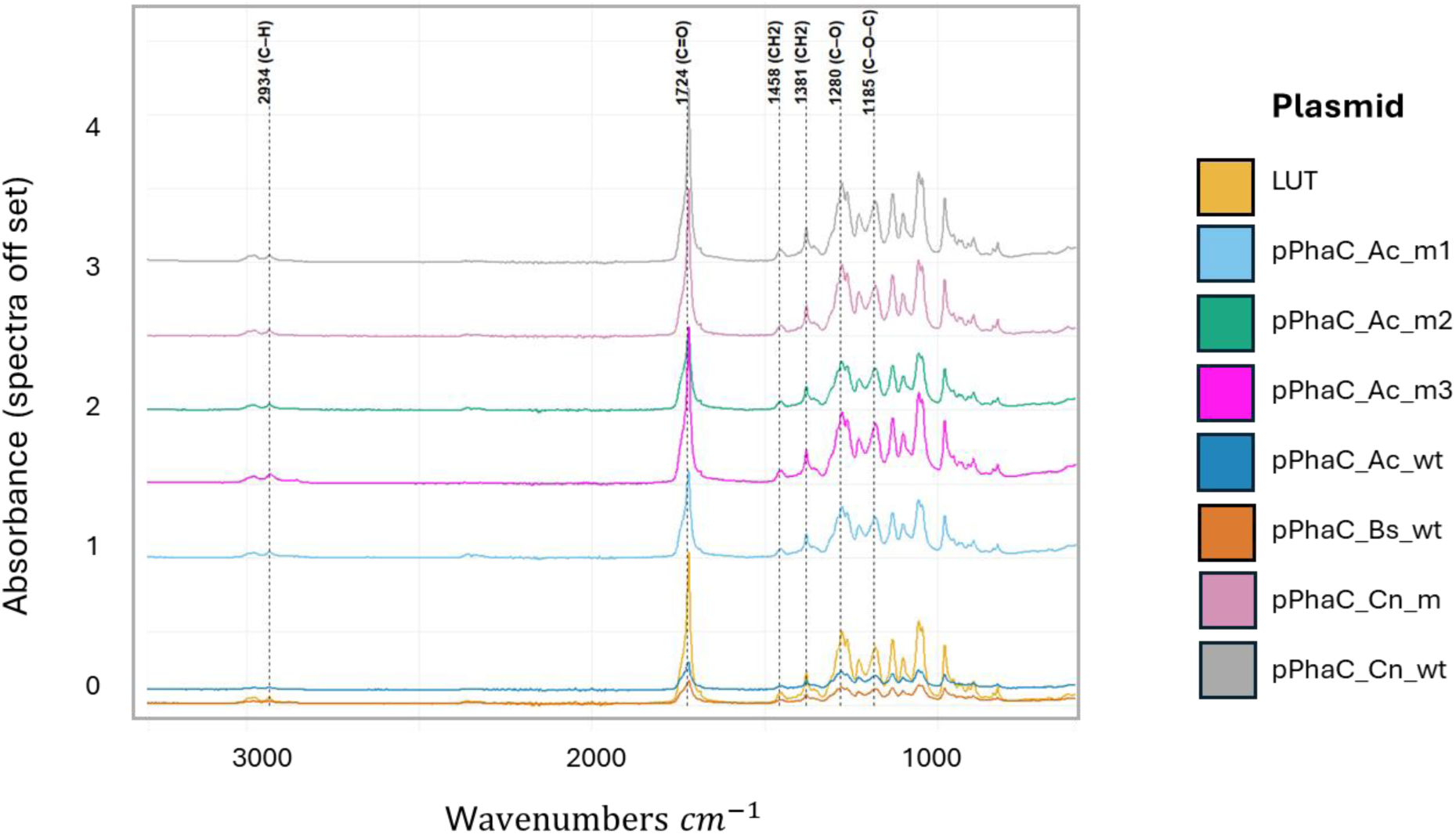
FTIR absorption spectra for PHA molecules from different strains with the Pha- variant library. The spectra show how different functional groups and molecular structures within these samples absorb infrared light at specific wavelengths, which can be linked to particular chemical bonds and structures. Most absorption spectra correspond to the C. necator control. The C=O stretching (peak around 1720 cm^-1^) that is observably wider in most of the transgenic PhaC enzyme variants, indicate an increase in co-3HHx (3-hydroxyhexanoate) portion of the P(3HB-co-3HHx) copolymers in comparison to the standard (LUT).

The asymmetric CH_3_ groups (*54*), are the side chains (R-group) of the repeating monomer unit. Particularly in the most common type P(3HB), where the R group is a methyl group attached to the third carbon (C3) of the hydroxyalkanoate backbone. This methyl side chain gives PHA its specific properties, and influences the crystallinity, melting point, and mechanical strength of the polymer (*55*, *56*). (*50*) Collectively, these peaks show the presence of ester groups and aliphatic chains, which are key structural features of PHAs. Other peaks relate to the hydrocarbon backbone and methyl groups of the polymer structure, typical for PHAs produced by microorganisms (*57*).

Differential scanning calorimetry (DSC) was used to characterize the thermal behaviour of PHA by measuring the glass transition temperature (T_g_), melting temperature (T_m_), and any cold crystallization events. These parameters describe crystallinity, chain mobility, and thermal stability of the material, with PHA from *C. necator* typically showing a sharp melting endotherm around 170–180 °C, sometimes with a double peak due to crystal thickness heterogeneity (*58*). In contrast, *A. caviae* PHA synthase often produces more flexible copolymers such as P(3HB-co-3HHx), where longer side-chain monomers like 3-hydroxyhexanoate lower crystallinity, depress and broaden T_m_ into the 120–160 °C range, and can reduce T_g_ below 0 °C (*59*).

In this study, T_g_ and T_m_ values for all variants fell within the literature ranges for P(3HB) and related copolymers, and the observed differences were small. PHA samples from pPhaC_Ac_wt and pPhaC_Bs_wt showed no clear T_g_, consistent with a more amorphous, less crystalline character. The small T_g_ increase in the *C. necator* mutant relative to wild type (8.34 °C vs 7.39 °C) is likely within experimental variation, whereas the lower T_g_ of the *A. caviae* double mutant (4.64 °C) agrees with the previously observed increased flexibility in *A. caviae* -derived copolymers (*60*). T_m_ values clustered around 172–174 °C, with an only slightly lower T_m_ for the *C. necator* variants, indicating small changes in crystallinity that are minor compared with the strong T_m_ depression seen for more comonomer--rich materials (Tab. 2) (*61*). Together, the different PhaC variants shift PHA thermal properties in the expected direction, but the resulting changes in T_g_ and T_m_ are generally small and fall within the range of DSC measurement uncertainty, as have been reported to vary between ∼0.5-5 °C (*62*, *63*).

**Table 2.**
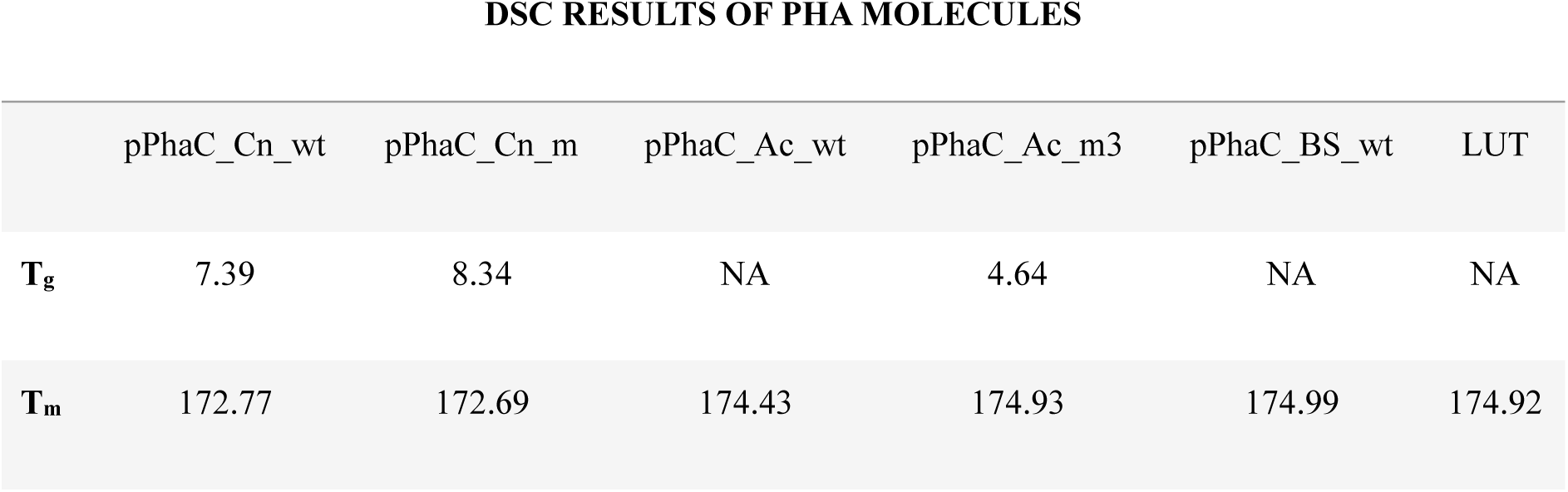
Differential Scanning Calorimetry (DSC) properties of PHA produced by different variants of the PhaC library. C. necator PhaC produced PHA with higher T_g_ and lower T_m_,,though variation falls within expected experimental uncertainty and hence does not suggest significant changes in flexibility or thermal stability..

Despite PhaCs catalysing the same reaction, our results show that the produced PHA polymers differ in their molecular structure, the quantity they are produced in and how they aggregate within the cell. A summary of the results is presented in Table 3. The PhaC variant from *A. caviae* with the double mutation appears to have the highest PHA yields that come in large granules in combination with a larger window between T_g_ and T_m_, while the wild-type version of this variant also achieves high yields in combination with an increased proportion of PHA positive cells.

**Table 3.**
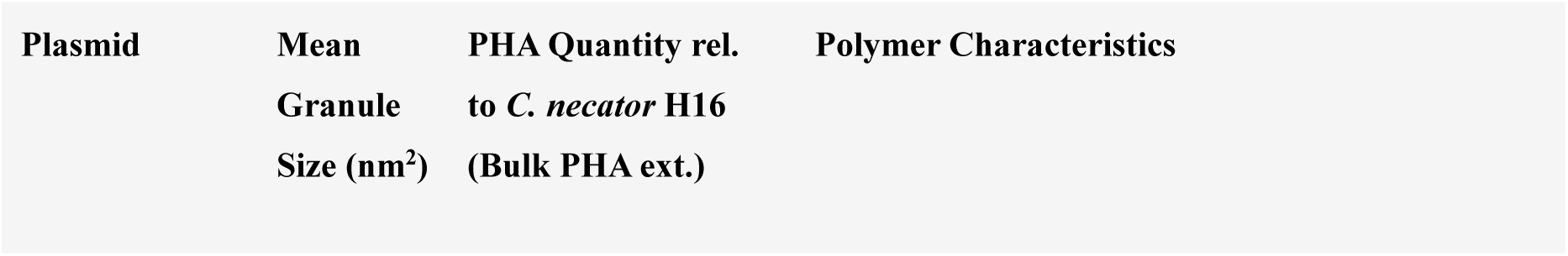
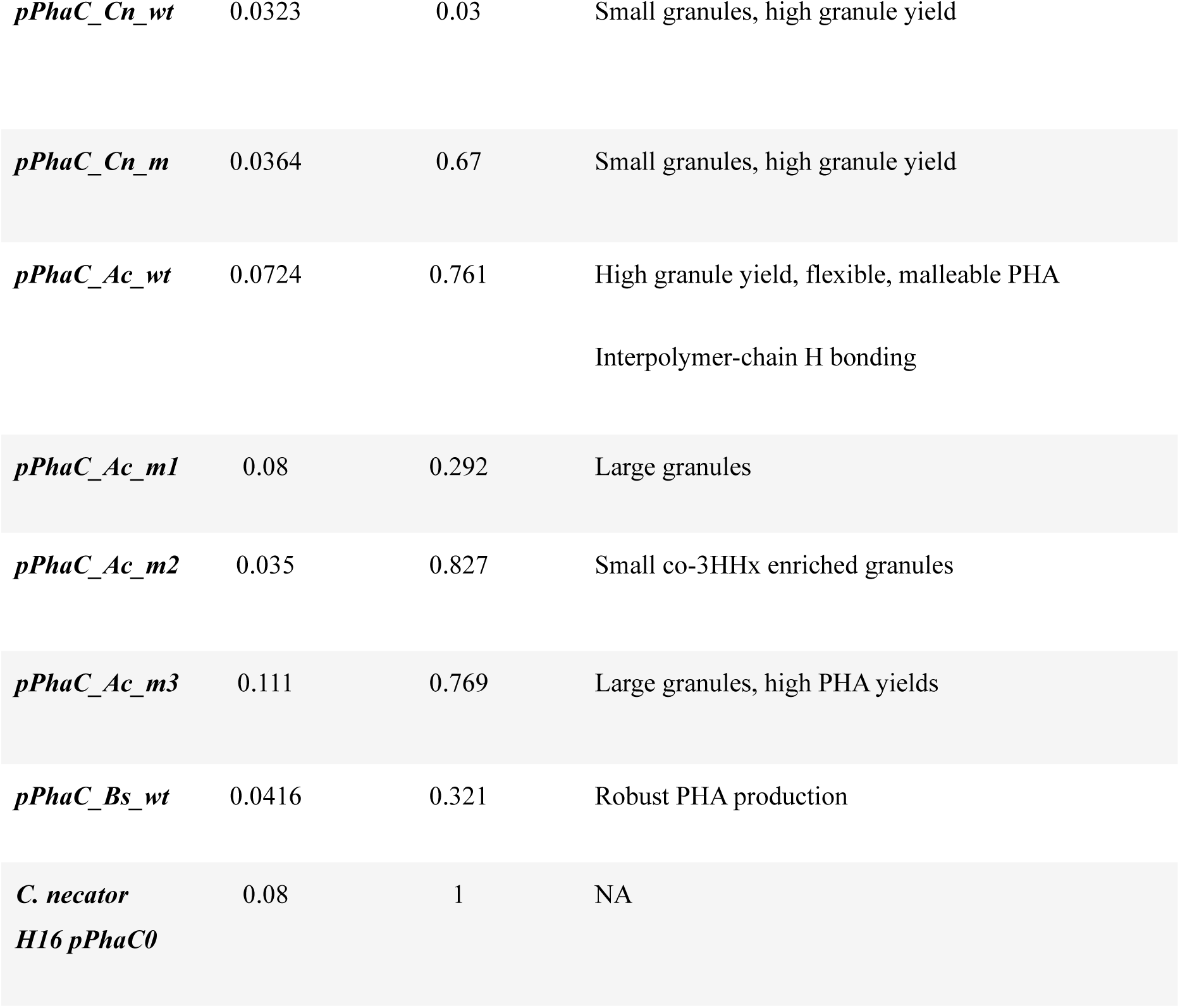
Summary of PHA production and PHA Granule properties by PhaC variant based on TEM, FTIR & DSC data.

### Co-culture system for enhanced PHA production

Having demonstrated the production of bio-PHA with *C. necator* is possible and polymer composition and granule size can be influenced by selection of the PhaC variant, we aimed to further optimize this biomanufacturing process by expanding the space of possible substrates for PHA production. Previously, Bhatia *et al.* (*64*) demonstrated that co-cultures of *C. necator* and *B. subtilis* were able to produce PHA with sucrose as sole carbon source. Usually, *C. necator* would not be able to use sucrose as substrate, however, *B. subtilis* breaks down sucrose into glucose and fructose. While glucose is directly used by *B. subtilis*, the fructose could then be used by *C. necator* to produce PHA. This is an attractive bioprocess because sucrose is an abundant feedstock, often present in waste products from processing of plant derived sugars, *e.g.* sugarcane molasses (*65*). However, achieving sustained and continuous production from synthetic microbial consortia and maintaining optimal concentrations of each strain in the co-culture is an ongoing endeavour.

When considering synthetic microbial consortia for bioproduction one must manage and control co-culture composition in a way that optimises specific bioproduction, in this case of PHA. To ensure sufficient substrate streams from *B. subtilis* to *C. necator*, we hypothesized that a higher *B. subtilis* percentage in the co-culture would be beneficial for PHA production. This led us to consider how a combination of controlled introduction of antibiotics and optimisation of the inoculation ratio in batch cultures might improve product yield. To this end, *B. subtilis* is significantly more tolerant to tetracycline than *C. necator*, which allows the control of the *C. necator* population. To investigate this dependence, we screened batch cultures with varying tetracycline concentration and inoculation ratios of *B. subtilis* (Figure 5). The maximum antibiotic concentration used was the lowest concentration at which no growth of *C. necator* was observed in previous experiments. In total, three tetracycline concentrations were tested, 0, 0.2 and 0.4 µg/mL.

**Figure 5.**
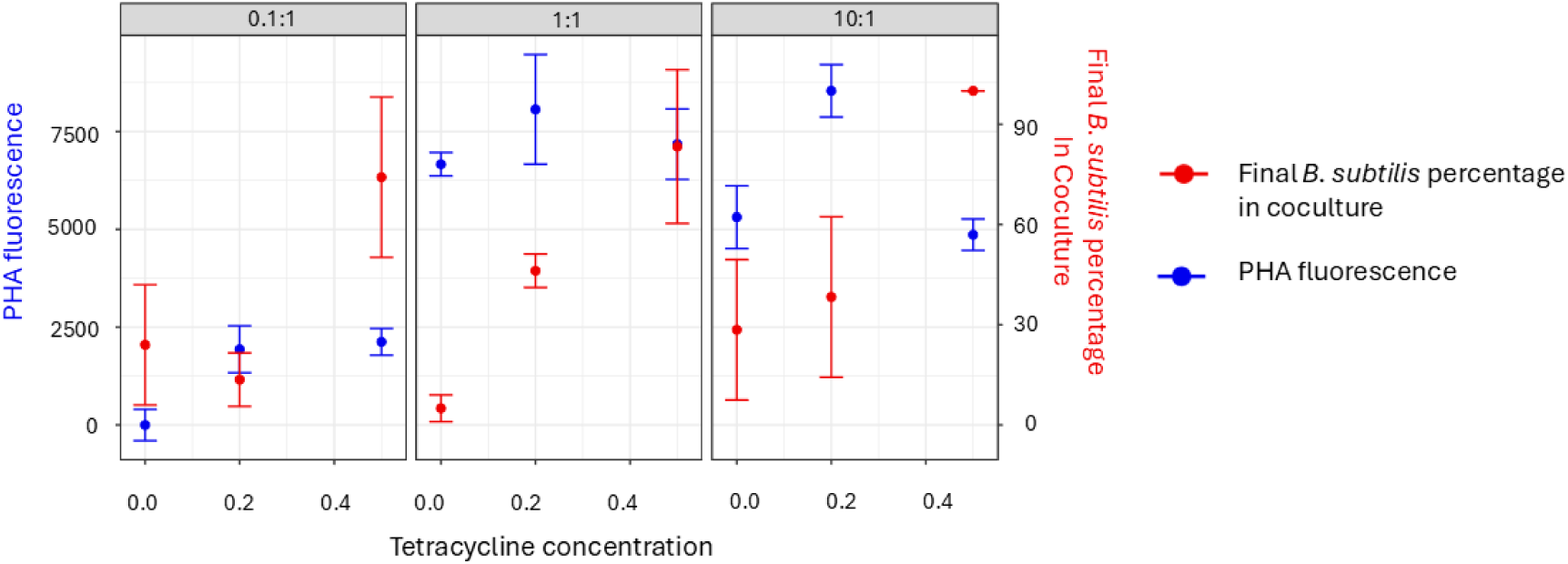
PHA production in C. necator in response to different co-culture parameters. The PHA production and final bacterial composition of the co-culture is dependent on B. subtills inoculation ratios and Tetracycline (µg/mL) concentrations. Maximal PHA production was achieved when Tetracycline was used in low concentrations (0.2 µg/ mL) to control the C. necator portion of the co-culture, as B. subtilis is more tolerant to this antibiotic than the PHA producing C. necator. The starting inoculation ratio of 0.1:1 showed the highest PHA production under all Tetracycline concentrations during this experiment, but the B. subtilis inoculation ratio of 10:1 in combination with the 0.2 µg/ ml antibiotics treatment achieved the highest total observed PHA production. Overall, manipulating co-culture composition and control proofed to be a beneficial strategy to maximise PHA production.

We observed that PHA production was strongly influenced by the *B. subtilis* inoculation ratio. Low starting ratios (0.1:1) resulted in minimal yields, whereas balanced or higher ratios supported greater production. At a 10:1 ratio, the highest yield occurred at 0.2 µg/mL tetracycline, while both, the absence and excess of antibiotics reduced PHA output due to population imbalance between *C. necator* and *B. subtilis*. At inoculation ratios of 1:1 and 10:1, the proportion of *B. subtilis* in the co-culture increased with tetracycline concentration (Figure 5).

Consequently, finding the right balance of the two co-culture species is crucial to maximise PHA production.

### Added functionality of PHA Bioplastic granules with SpyTag-SpyCatcher system

With the ability to use *C. necator* as a microbial cell factory to produce PHA granules of tuneable size and material properties, we developed this further to add bespoke functions to these bio-nanoparticles. We used the SpyTag-SpyCatcher system, which functions as a peptide-protein pair that forms irreversible covalent bounds when mixed. Adding the SpyCatcher (protein) part of the system to one component and the SpyTag (peptide) to another will lead to efficient binding of the tagged compound together with the catcher (*66*, *67*).

To attach the system to our PHA granules we leveraged the fact that the PhaC enzyme is covalently attached to nascent PHA polymer chains and remains attached to the granule surface at the terminating end of the polymer (*68*, *69*). By adding an N-terminal translational fusion of the SpyTag003 sequence to the *phaC* coding region of pPhaC_Ac_m1, we produced pPhaC_Ac_m1::SpyTag fusions. Granules produced with this new enzyme should bear the SpyTag sequence at their surface, such that proteins bearing the SpyCatcher domain can bind to them specifically and permanently.

As proof of concept, we used *E. coli* to produce GFP_SC003, a GFP protein variant bearing a C-terminal SpyCatcher003 domain, and *C. necator ΔphaC* bearing the pPhaC_Ac_m1::SpyTag plasmid to produce SpyTag bearing granules (Fig. 6 A).

**Figure 6.**
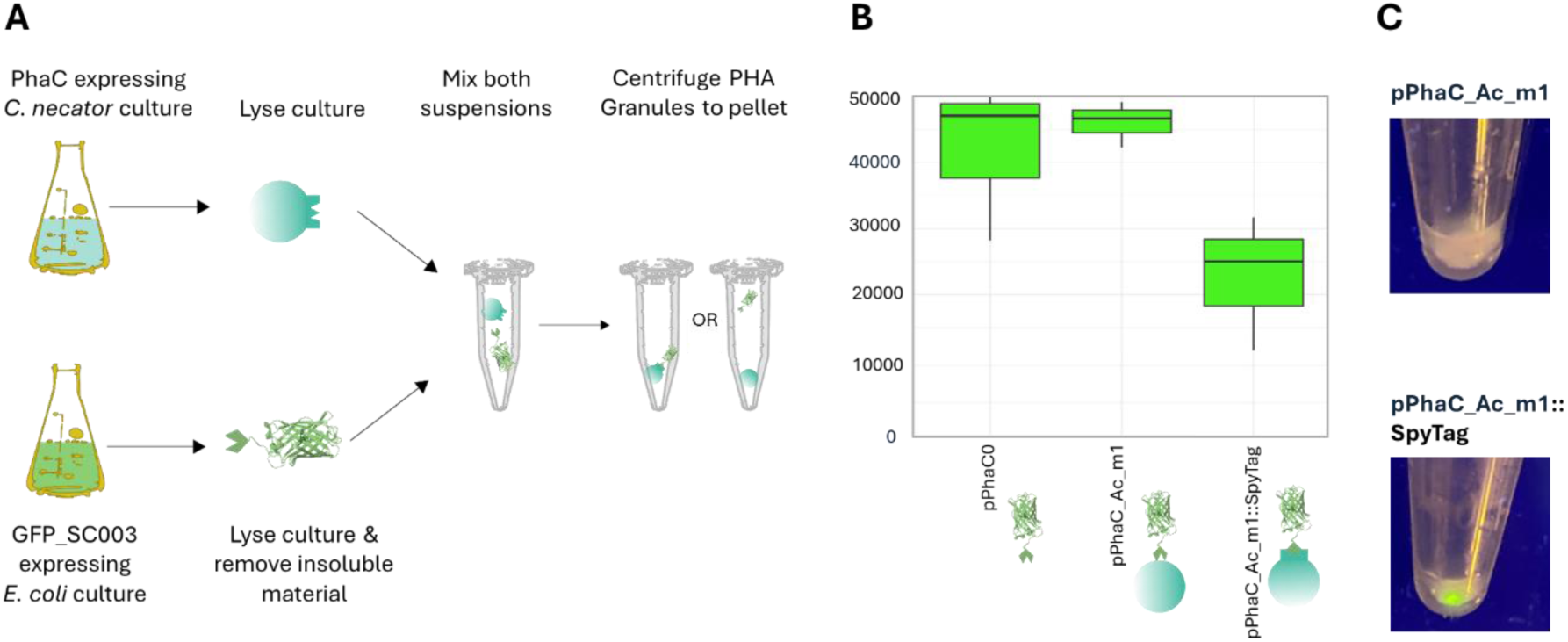
Functionalisation of PHA Granule Nanoparticles. GFP accumulation as example. **A)** Separate cultures of PHA-SpyTag producing C. necator and E. coli cultures producing GFP_SpyCatch003 (SC003) were generated. Cultures were lysed by sonication and E. coli cultures were centrifuged to remove insoluble residues. C. necator lysates were normalised to match A280 for protein content and mixed independently with identical suspensions of E. coli lysate containing GFP_SC003. After approximately 2 hours of room temperature incubation, cultures were centrifuged to separate soluble GFP_SC003 from that which may have been attached to granules. **B)** The remaining GFP in the soluble fraction was measured using a plate reader. Both control plasmids, pPhaC0 and pPhaC_Ac_m1 had high GFP concentrations in the supernatant, while the SpyTag version reduced the GFP content to almost the half in the supernatant by pelleting it with the pPhaC_Ac_m1::SpyTag nanoparticles. **C)** Pellets were imaged under a 450nm blue LED light, showing accumulation of GFP.

To test the system, lysates of these cultures were mixed and insoluble granules separated by centrifugation (see Methods). GFP fluorescence was measured in the supernatants, with the hypothesis that if GFP has bound to granules, it should be removed with the pellet and fluorescence in the liquid fraction should be reduced. pPhaC_Ac_m1 was used as a negative control, lacking the SpyTag003 sequence, and pPHAC0 was used as a control for any non-specific binding. As expected, GFP fluorescence was reduced when the pPhaC_Ac_m1::SpyTag plasmid was used (Fig. 6 B), and GFP could be observed by eye in the pellet containing the PHA granules (Fig. 6 C). The specific version of the SpyTag-SpyCatcher we used in this study was chosen for its very high affinity (*70*) and has been deployed efficiently in various settings before from *in vivo* physiological conditions (*71*) to high pH (*72*) environments, indicating they work reliably and efficiently in most settings.

## Conclusion

This work establishes *Cupriavidus necator* as a versatile microbial chassis for the production of tuneable, functionalised PHA nanoparticles by integrating enzyme engineering, materials characterisation and co-culture process control. We demonstrated that the transformation efficiency of *C. necator* can effectively be optimized by increasing the OD when preparing competent cells, using low DNA concentrations and choosing an optimal field strength and outgrowth time. By constructing and screening a phaC variant library, we showed that polymer yield, granule morphology and thermal properties can be rationally adjusted to generate materials spanning from highly crystalline PHAs to softer copolymers with enlarged processing windows. Coupling *C. necator* with *B. subtilis* in antibiotic-tuned co-cultures further broadens accessible feedstocks and significantly enhances PHA production, while SpyTag-SpyCatcher functionalisation demonstrates that granules can act as modular scaffolds for protein display and cargo capture. This conceptual framework enables a broad range of new nanotechnology applications. By integrating the SpyTag-SpyCatch system with the *C. necator* - *B. subtilis* co-culture, the entire process could be streamlined into a one-pot synthesis in which both polymer production and functionalisation occur in the same bioreactor. For example, one strain could be engineered to produce PHA granules displaying SpyTag, while the other expresses SpyCatch-fused enzymes, fluorophores, or targeting ligands that covalently attach to the polymer surface. Building a modular “toolbox” of orthogonal tag-catch pairs would then allow different strains and functions to be mixed and matched without re-engineering the core system each time. In practice, this might mean swapping in a SpyPair that recruits metal-binding peptides for biosensing, another that immobilises redox enzymes for bioelectrocatalysis, or one that displays cell-targeting peptides for drug delivery, all using the same underlying co-culture platform. Together, these advances provide a toolbox that moves bioPHA from a bulk plastic substitute toward a platform of high-value designer bionanoparticles with possible applications in sustainable biomanufacturing, biomedical applications and future bioremediation technologies.

## Methods

### Strains and media conditions

Strains used were *B. subtilis* 168 and *C. necator* str. H16. These were cultured routinely in Lysogeny Broth (LB, Formedium) for starter cultures and solid media. For PHA production by *C. necator* and derivative strains, Low Nitrogen M9 was used (Formedium). For co-culture experiments in a plate reader, a Tecan Spark plate reader was used, and media was composed of 5x M9 (Formedium), Trace Elements, 50 µg/mL Tryptone, 0.4 % (m/v) sucrose, 50 µM CaCl_2_ and 2 mM MgSO_4_. For turbidostat co-cultures, an identical medium was used with the addition of 0.025% (m/v) Cas-amino acids (Formedium). As selection marker for the transgenic plasmids the antibiotic kanamycin was added at a final concentration of 100 µg/mL. For inducible constructs, 2 mM arabinose were added to the media.

### Cloning

Synthetic DNA encoding the transgenic phaC sequences were ordered from IDT as gBlocks. These were assembled into the pRH412 backbone (a kind gift of Robert Habgood) using the NEB HiFi DNA Assembly kit. Primers and plasmids are available in the Supplementary Section S2 and S3. PCR was performed using the Q5 DNA polymerase. To introduce mutations, PCR-based plasmid mutagenesis was used to produce phaC allele variants. The KLD Enzyme kit by NEB was used to perform the final assemblies. Primers used are available in Supplementary Material S2. DH5α *E. coli* was used for all routine cloning procedures.

### Plasmid Delivery by Electroporation of *C. necator*

Electrocompetent *C. necator* cells were prepared by adapting a protocol established for *Pseudomonas aeruginosa* (*73*). Briefly, cells from LB plates grown for 48h were cultured sequentially in SOB medium to the desired OD (0.2–5), washed twice in 1 mM MgSO₄, resuspended in 25% glycerol, aliquoted (50 µl), and stored at −80 °C. For transformation, thawed competent cells were mixed with 0–500 ng plasmid DNA and electroporated using 0.1–0.2 cm cuvettes at 1.8–2.5 kV on a Bio-Rad MicroPulser (Ec1 or Ec2 setting). Following a 2–4 h recovery in SOB at 30 °C, cultures were diluted and plated on selective or non-selective LB media to determine transformation efficiency (CFU/µg DNA) as shown in Figure S1.

### PHA estimation by Fluorescence and Flow Cytometry

Cultures were grown in 5 mL of Low Nitrogen M9 medium (see above) with arabinose in Falcon tubes with orbital shaking at 250 rpm for approximately 16 hours. Cells were washed in sterile PBS and then mixed with an equal volume of 0.02% (m/v) Nile Blue A (Sigma). Cells were incubated for at least three minutes before fluorescence was measured with a Tecan Spark plate reader at 450 nm Excitation and 550 nm Emission. For flow cytometery, a BD FACS Aria III flow cytometer was used, with a 488 nm laser. Flowing Software (*74*) was used to analyse data held in resulting .fcs files. For the gating strategy to identify PHA positive cells, singlet cell events were first identified and gated based on forward (FSC) and side scatter (SSC) to exclude debris and aggregates. A fluorescence gate for PHA-positive cells was then defined on the Nile Blue A emission channel using an unstained control to set the fluorescence threshold. The percentage of PHA-positive cells was calculated as the number of events within the Nile Blue A-positive gate divided by the total number of gated singlet cells, multiplied by 100.

### TEM and PHA Granule Morphology Characterisation

Cells were cultured identically as for Flow Cytometry above. Cells were washed in PBS before the pellet was fixed in fixing buffer which contained 1 mL 25% glutaraldehyde, 2.5 mL 16% paraformaldehyde, 5 mL 0.2 M sodium cacodylate, and 1.5 mL distilled water. Following this, samples were processed by the Electron Microscopy Facility at the Sir William Dunn School of Pathology, University of Oxford and imaged using a JEOL 1400 TEM.

PHA granules inside cells were analysed using the Fiji application for ImageJ. A Weka Trainable Image Classifier was deployed on normalised raw images (.dm4 file format) to build an image classifier which predicts probability of specific pixels representing cells, granules or background. The SlideSet plugin (*75*) was used to apply this classifier and obtain probability maps for each .dm4 file. These were then processed by enhancing the contrast and normalising histograms in the resulting binary images. This produces an image with pixels coloured between 0 (black) and 255 (white), where ‘whiter’ pixels indicate a greater confidence that that pixel represents a PHA granule. A threshold was set to 175 contrast for each image to produce a binary image of only black (0) or white pixels (255). Next, Particle Analysis in ImageJ was used to identify PHA granules, with constraints of circularity between 0.12 and 1 and area of particle of at least 0.01 square microns.

### PHA Purification and Materials Analysis

*C. necator* cells bearing plasmids specified were cultured in starter cultures in LB with kanamycin overnight. These were then inoculated into 100 mL Low Nitrogen M9 with the addition of arabinose and kanamycin for approximately 24 hours. Cells were then harvested by centrifugation and resuspended in 10 mL 30 % (v/v) sodium hypochlorite. These were incubated for an hour on ice, before 5 mL distilled water was added and the suspension vortexed thoroughly. Bulk PHA was extracted as the insoluble fraction by centrifugation. This was washed three times with distilled water, then two to three times with methanol. The purified PHA was then air dried by leaving the caps of the tubes off.

Samples were analysed by FTIR and DSC by Oxford Materials Characterization Service. The DSC was run on a Perkin Elmer Hyper DSC under nitrogen and the FTIR was run on a Shimadzu IRSpirit FTIR with a QATR-S Single Bounce Diamond ATR (Attenuated Internal Reflection) sampling set up.

### Turbidostat Co-culture

Starter cultures in LB were prepared of *B. subtilis* and *C. necator* by inoculating a single colony from a streak on an LB agar plate into 5 mL LB liquid media and incubating at 30 °C with shaking overnight. Cultures were then washed in M9 medium as described above and diluted to an optical density of 0.3 prior to commencing the experiment. At OD 0.3 *C. necator* suspensions contain approximately 10x more CFU/mL than *B. subtilis* at the same OD600. As such, a suspension containing approximately equal concentrations of each strain was prepared. 200 µL of this suspension was inoculated into Chi.Bio bioreactors (*76*). Cultures were grown in turbidostat mode with a target OD600 of 0.3 in M9 as described above at 30 °C and stirring set to 0.5. Initially, cultures were left in unmodified M9, however after approximately 24 hours, Tetracycline was added at concentrations specified. This was repeated every 24 hours for as long as the experiment proceeded. At each of these moments, a 0.5 mL sample was also obtained and mixed with 0.5 mL 50% (v/v) glycerol and flash-frozen. Culture composition was probed by thawing these samples and serially diluting them 10-fold eventually reaching a dilution factor of 10^-8^ of the original sample concentration. 5 µL of each dilution was plated on LB agar and incubated overnight at 30 °C. After this point, the faster growing *B. subtilis* colonies could be counted. After a subsequent overnight incubation, the *C. necator* colonies had formed and could be counted.

## Acknowledgements

The authors recognised support from Engineering and Physical Sciences Research Council (EPSRC) project EP/W000326/1.

## References

1. Md. G. Kibria, N. I. Masuk, R. Safayet, H. Q. Nguyen, M. Mourshed, Plastic Waste: Challenges and Opportunities to Mitigate Pollution and Effective Management. Int. J. Environ. Res. 17, 20 (2023).

2. F. Bauer, T. D. Nielsen, L. J. Nilsson, E. Palm, K. Ericsson, A. Fråne, J. Cullen, Plastics and climate change—Breaking carbon lock-ins through three mitigation pathways. One Earth 5, 361–376 (2022).

3. K. Houssini, J. Li, Q. Tan, Complexities of the global plastics supply chain revealed in a trade-linked material flow analysis. Commun. Earth Environ. 6, 257 (2025).

4. E. Van Roijen, S. A. Miller, Leveraging biogenic resources to achieve global plastic decarbonization by 2050. Nat. Commun. 16, 7659 (2025).

5. S. Pratt, L.-J. Vandi, D. Gapes, A. Werker, A. Oehmen, B. Laycock, “Polyhydroxyalkanoate (PHA) Bioplastics from Organic Waste” in Biorefinery: Integrated Sustainable Processes for Biomass Conversion to Biomaterials, Biofuels, and Fertilizers, J.-R. Bastidas-Oyanedel, J. E. Schmidt, Eds. (Springer International Publishing, Cham, 2019; 10.1007/978-3-030-10961-5_26), pp. 615–638.

6. G. Atiwesh, A. Mikhael, C. C. Parrish, J. Banoub, T.-A. T. Le, Environmental impact of bioplastic use: A review. Heliyon 7 (2021).

7. B. Dalton, P. Bhagabati, J. De Micco, R. B. Padamati, K. O’Connor, A Review on Biological Synthesis of the Biodegradable Polymers Polyhydroxyalkanoates and the Development of Multiple Applications. Catalysts 12, 319 (2022).

8. R. Ben Abdeladhim, J. A. Reis, A. M. Vieira, C. D. de Almeida, Polyhydroxyalkanoates: Medical Applications and Potential for Use in Dentistry. Materials 17, 5415 (2024).

9. D. A. Gregory, C. S. Taylor, A. T. R. Fricker, E. Asare, S. S. V. Tetali, J. W. Haycock, I. Roy, Polyhydroxyalkanoates and their advances for biomedical applications. Trends Mol. Med. 28, 331–342 (2022).

10. S. Ray, V. C. Kalia, Biomedical Applications of Polyhydroxyalkanoates. Indian J. Microbiol. 57, 261–269 (2017).

11. K. Verma, C. Sarkar, S. Saha, Exploration of biodegradable polymeric particles in agriculture: a holistic approach for sustainable farming. *Environ*. Sci. Adv. 4, 409–431 (2025).

12. E. P. Lamparelli, M. Marino, M. A. Szychlinska, N. Della Rocca, M. C. Ciardulli, P. Scala, R. D’Auria, A. Testa, A. Viggiano, F. Cappello, R. Meccariello, G. Della Porta, A. Santoro, The Other Side of Plastics: Bioplastic-Based Nanoparticles for Drug Delivery Systems in the Brain. Pharmaceutics 15, 2549 (2023).

13. R. Sreena, A. J. Nathanael, Biodegradable Biopolymeric Nanoparticles for Biomedical Applications-Challenges and Future Outlook. Materials 16, 2364 (2023).

14. J. Panich, B. Fong, S. W. Singer, Metabolic Engineering of *Cupriavidus necator* H16 for Sustainable Biofuels from CO2. Trends Biotechnol. 39, 412–424 (2021).

15. L. Zhang, Z. Jiang, T.-H. Tsui, K.-C. Loh, Y. Dai, Y. W. Tong, A Review on Enhancing Cupriavidus necator Fermentation for Poly(3-hydroxybutyrate) (PHB) Production From Low-Cost Carbon Sources. Front. Bioeng. Biotechnol. 10, 946085 (2022).

16. M. Weldon, C. Euler, Physiology-informed use of Cupriavidus necator in biomanufacturing: a review of advances and challenges. Microb. Cell Factories 24, 30 (2025).

17. A. Zúñiga, F. de la Fuente, F. Federici, C. Lionne, J. Bônnet, V. de Lorenzo, B. González, An Engineered Device for Indoleacetic Acid Production under Quorum Sensing Signals Enables Cupriavidus pinatubonensis JMP134 To Stimulate Plant Growth. ACS Synth. Biol. 7, 1519–1527 (2018).

18. S. Mishra, P. M. Perkovich, W. P. Mitchell, M. Venkataraman, B. F. Pfleger, Expanding the synthetic biology toolbox of Cupriavidus necator for establishing fatty acid production. J. Ind. Microbiol. Biotechnol. 51 (2024).

19. R. K. Purama, J. N. Al-Sabahi, K. Sudesh, Evaluation of date seed oil and date molasses as novel carbon sources for the production of poly(3Hydroxybutyrate-*co*-3Hydroxyhexanoate) by *Cupriavidus necator* H16 Re 2058/pCB113. Ind. Crops Prod. 119, 83–92 (2018).

20. R. R. Dalsasso, F. A. Pavan, S. E. Bordignon, G. M. F. de Aragão, P. Poletto, Polyhydroxybutyrate (PHB) production by *Cupriavidus necator* from sugarcane vinasse and molasses as mixed substrate. Process Biochem. 85, 12–18 (2019).

21. M. S. Morlino, R. Serna García, F. Savio, G. Zampieri, T. Morosinotto, L. Treu, S. Campanaro, *Cupriavidus necator* as a platform for polyhydroxyalkanoate production: An overview of strains, metabolism, and modeling approaches. Biotechnol. Adv. 69, 108264 (2023).

22. M. Koch, K. Forchhammer, Polyhydroxybutyrate: A Useful Product of Chlorotic Cyanobacteria. Microb. Physiol. 31, 67–77 (2021).

23. R. Mozes-Koch, E. Tanne, A. Brodezki, R. Yehuda, O. Gover, H. D. Rabinowitch, I. Sela, Expression of the entire polyhydroxybutyrate operon of Ralstonia eutropha in plants. J. Biol. Eng. 11, 44 (2017).

24. D. Jendrossek, Polyhydroxyalkanoate Granules Are Complex Subcellular Organelles (Carbonosomes). J. Bacteriol. 191, 3195–3202 (2009).

25. S. Zher Neoh, M. Fey Chek, H. Tiang Tan, J. A. Linares-Pastén, A. Nandakumar, T. Hakoshima, K. Sudesh, Polyhydroxyalkanoate synthase (PhaC): The key enzyme for biopolyester synthesis. Curr. Res. Biotechnol. 4, 87–101 (2022).

26. K. Jia, R. Cao, D. H. Hua, P. Li, Study of Class I and Class III Polyhydroxyalkanoate (PHA) Synthases with Substrates Containing a Modified Side Chain. Biomacromolecules 17, 1477–1485 (2016).

27. H. J. Tang, S. Z. Neoh, K. Sudesh, A review on poly(3-hydroxybutyrate-co-3-hydroxyhexanoate) [P(3HB-co-3HHx)] and genetic modifications that affect its production. Front. Bioeng. Biotechnol. 10 (2022).

28. B. H. A. Rehm, Polyester synthases: natural catalysts for plastics. Biochem. J. 376, 15–33 (2003).

29. F. A. El-malek, A. Steinbüchel, Post-Synthetic Enzymatic and Chemical Modifications for Novel Sustainable Polyesters. Front. Bioeng. Biotechnol. 9 (2022).

30. M. F. Chek, S.-Y. Kim, T. Mori, H. Arsad, M. R. Samian, K. Sudesh, T. Hakoshima, Structure of polyhydroxyalkanoate (PHA) synthase PhaC from Chromobacterium sp. USM2, producing biodegradable plastics. Sci. Rep. 7, 5312 (2017).

31. C. F. Budde, S. L. Riedel, L. B. Willis, C. Rha, A. J. Sinskey, Production of Poly(3-Hydroxybutyrate-co-3-Hydroxyhexanoate) from Plant Oil by Engineered Ralstonia eutropha Strains▿. Appl. Environ. Microbiol. 77, 2847–2854 (2011).

32. K. Bhubalan, D.-N. Rathi, H. Abe, T. Iwata, K. Sudesh, Improved synthesis of P(3HB-*co*-3HV-*co*-3HHx) terpolymers by mutant *Cupriavidus necator* using the PHA synthase gene of *Chromobacterium* sp. USM2 with high affinity towards 3HV. Polym. Degrad. Stab. 95, 1436–1442 (2010).

33. K. Harada, S. Kobayashi, K. Oshima, S. Yoshida, T. Tsuge, S. Sato, Engineering of Aeromonas caviae Polyhydroxyalkanoate Synthase Through Site-Directed Mutagenesis for Enhanced Polymerization of the 3-Hydroxyhexanoate Unit. Front. Bioeng. Biotechnol. 9, 627082 (2021).

34. V. Peters, B. H. A. Rehm, In Vivo Enzyme Immobilization by Use of Engineered Polyhydroxyalkanoate Synthase. Appl. Environ. Microbiol. 72, 1777–1783 (2006).

35. V. Peters, B. H. A. Rehm, In vivo monitoring of PHA granule formation using GFP-labeled PHA synthases. FEMS Microbiol. Lett. 248, 93–100 (2005).

36. K. L. Tee, J. Grinham, A. M. Othusitse, M. González-Villanueva, A. O. Johnson, T. S. Wong, An Efficient Transformation Method for the Bioplastic-Producing “Knallgas” Bacterium Ralstonia eutropha H16. Biotechnol. J. 12, 1700081 (2017).

37. S. Arhar, T. Rauter, H. Stolterfoht-Stock, V. Lambauer, R. Kratzer, M. Winkler, M. Karava, R. Kourist, A. Emmerstorfer-Augustin, CO2-based production of phytase from highly stable expression plasmids in Cupriavidus necator H16. Microb. Cell Factories 23, 9 (2024).

38. T. P. Huang, Z. J. Heins, S. M. Miller, B. G. Wong, P. A. Balivada, T. Wang, A. S. Khalil, D. R. Liu, High-throughput continuous evolution of compact Cas9 variants targeting single-nucleotide-pyrimidine PAMs. Nat. Biotechnol. 41, 96–107 (2023).

39. A. Khlebnikov, Ø. Risa, T. Skaug, T. A. Carrier, J. D. Keasling, Regulatable Arabinose-Inducible Gene Expression System with Consistent Control in All Cells of a Culture. J. Bacteriol. 182, 7029–7034 (2000).

40. G. T. Little, M. Ehsaan, C. Arenas-López, K. Jawed, K. Winzer, K. Kovacs, N. P. Minton, Complete Genome Sequence of Cupriavidus necator H16 (DSM 428). Microbiol. Resour. Announc. 8, 10.1128/mra.00814-19 (2019).

41. S. Taguchi, H. Nakamura, T. Hiraishi, I. Yamato, Y. Doi, In Vitro Evolution of a Polyhydroxybutyrate Synthase by Intragenic Suppression-Type Mutagenesis. J. Biochem. (Tokyo) 131, 801–806 (2002).

42. E. Shimamura, K. Kasuya, G. Kobayashi, T. Shiotani, Y. Shima, Y. Doi, Physical Properties and Biodegradability of Microbial Poly(3-hydroxybutyrate-co-3-hydroxyhexanoate), ACS Publications (2002). 10.1021/ma00081a041.

43. T. Kichise, S. Taguchi, Y. Doi, Enhanced accumulation and changed monomer composition in polyhydroxyalkanoate (PHA) copolyester by in vitro evolution of Aeromonas caviae PHA synthase. Appl. Environ. Microbiol. 68, 2411–2419 (2002).

44. T. Tsuge, Y. Saito, Y. Kikkawa, T. Hiraishi, Y. Doi, Biosynthesis and compositional regulation of poly[(3-hydroxybutyrate)-co-(3-hydroxyhexanoate)] in recombinant ralstonia eutropha expressing mutated polyhydroxyalkanoate synthase genes. Macromol. Biosci. 4, 238–242 (2004).

45. A. Amara, A. Steinbüchel, B. Rehm, In vivo evolution of the Aeromonas punctata polyhydroxyalkanoate (PHA) synthase: isolation and characterization of modified PHA synthases with enhanced activity. Appl. Microbiol. Biotechnol. 59, 477–482 (2002).

46. T. Tsuge, S. Watanabe, D. Shimada, H. Abe, Y. Doi, S. Taguchi, Combination of N149S and D171G mutations in Aeromonas caviae polyhydroxyalkanoate synthase and impact on polyhydroxyalkanoate biosynthesis. FEMS Microbiol. Lett. 277, 217–222 (2007).

47. N. G. Assefa, H. Hansen, B. Altermark, A unique class I polyhydroxyalkanoate synthase (PhaC) from Brevundimonas sp. KH11J01 exists as a functional trimer: A comparative study with PhaC from Cupriavidus necator H16. New Biotechnol. 70, 57–66 (2022).

48. S. Nakamura, Y. Doi, M. Scandola, Microbial synthesis and characterization of poly(3-hydroxybutyrate-co-4-hydroxybutyrate). Macromolecules 25, 4237–4241 (1992).

49. D. Jendrossek, Peculiarities of PHA granules preparation and PHA depolymerase activity determination. Appl. Microbiol. Biotechnol. 74, 1186–1196 (2007).

50. T. Van Nguyen, T. Nagata, K. Noso, K. Kaji, H. Masunaga, T. Hoshino, T. Hikima, S. Sakurai, K. Yamamoto, Y. Miura, T. Aoki, H. Yamane, S. Sasaki, Effect of the 3-Hydroxyhexanoate Content on Melt-Isothermal Crystallization Behavior of Microbial Poly(3-hydroxybutyrate-co-3-hydroxyhexanoate). Macromolecules 54, 8738–8750 (2021).

51. S. Bayarı, F. Severcan, FTIR study of biodegradable biopolymers: P(3HB), P(3HB-co-4HB) and P(3HB-co-3HV). J. Mol. Struct. 744–747, 529–534 (2005).

52. M. Kumar, A. Singhal, P. K. Verma, I. S. Thakur, Production and Characterization of Polyhydroxyalkanoate from Lignin Derivatives by Pandoraea sp. ISTKB. ACS Omega 2, 9156–9163 (2017).

53. C. Trakunjae, A. Boondaeng, W. Apiwatanapiwat, P. Janchai, S. Z. Neoh, K. Sudesh, P. Vaithanomsat, Statistical optimization of P(3HB-co-3HHx) copolymers production by Cupriavidus necator PHB−4/pBBR_CnPro-phaCRp and its properties characterization. Sci. Rep. 13, 9005 (2023).

54. C. R. Rech, S. M. Martelli, B. E. B. e Silva, K. C. da S. Brabes, Antimicrobial Analysis and Characterization of P(3HB) Films Containing Essential oils. *Orbital Electron*. J. Chem., 9–13 (2018).

55. A. Dhaini, V. Hardouin-Duparc, A. Alaaeddine, J.-F. Carpentier, S. M. Guillaume, Recent advances in polyhydroxyalkanoates degradation and chemical recycling. Prog. Polym. Sci. 149, 101781 (2024).

56. R. Sehgal, R. Gupta, Polyhydroxyalkanoate and its efficient production: an eco-friendly approach towards development. 3 Biotech 10,549 (2020).

57. S. Alfano, F. Pagnanelli, A. Martinelli, Rapid Estimation of Poly(3-hydroxybutyrate-co-3-hydroxyvalerate) Composition Using ATR-FTIR. Polymers 15, 4127 (2023).

58. N. Hernández-Herreros, A. Rodriguez, V. Rivero-Buceta, A. Rojas, M. A. Prieto, Flexible feeding strategy for high-yield PHA bioprocessing in *Cupriavidus necator* H16 from anaerobically fermented industrial wastewater. Bioresour. Technol. 434, 132774 (2025).

59. S. Y. Li, C. L. Dong, S. Y. Wang, H. M. Ye, G.-Q. Chen, Microbial production of polyhydroxyalkanoate block copolymer by recombinant Pseudomonas putida. Appl. Microbiol. Biotechnol. 90, 659–669 (2011).

60. P. K. Sharma, R. I. Munir, W. Blunt, C. Dartiailh, J. Cheng, T. C. Charles, D. B. Levin, Synthesis and Physical Properties of Polyhydroxyalkanoate Polymers with Different Monomer Compositions by Recombinant Pseudomonas putida LS46 Expressing a Novel PHA SYNTHASE (PhaC116) Enzyme. Appl. Sci. 7, 242 (2017).

61. H. J. Jung, B. Kim, T.-R. Choi, S. J. Oh, S. Kim, Y. Lee, Y. Shin, S. Choi, J. Oh, S. Y. Park, Y. S. Lee, Y. H. Choi, Y.-H. Yang, Novel differential scanning calorimetry (DSC) application to select polyhydroxyalkanoate (PHA) producers correlating 3-hydroxyhexanoate (3-HHx) monomer with melting enthalpy. Bioprocess Biosyst. Eng. 47, 1619–1631 (2024).

62. C. Dartiailh, W. Blunt, P. K. Sharma, S. Liu, N. Cicek, D. B. Levin, The Thermal and Mechanical Properties of Medium Chain-Length Polyhydroxyalkanoates Produced by Pseudomonas putida LS46 on Various Substrates. Front. Bioeng. Biotechnol. 8, 617489 (2021).

63. K. Grgurević, M. M. Nikolić, D. Kučić Grgić, V. O. Bulatović, Thermal, Mechanical, and Barrier Properties of PHBV Nanocomposites via TiO2 Incorporation for Sustainable Food Packaging. Polymers 18, 11 (2025).

64. S. K. Bhatia, J. J. Yoon, H. J. Kim, J. W. Hong, Y. Gi Hong, H. S. Song, Y. M. Moon, J. M. Jeon, Y. G. Kim, Y. H. Yang, Engineering of artificial microbial consortia of Ralstonia eutropha and Bacillus subtilis for poly(3-hydroxybutyrate-co-3-hydroxyvalerate) copolymer production from sugarcane sugar without precursor feeding. Bioresour. Technol. 257, 92–101 (2018).

65. R. R. Dalsasso, F. A. Pavan, S. E. Bordignon, G. M. F. de Aragão, P. Poletto, Polyhydroxybutyrate (PHB) production by Cupriavidus necator from sugarcane vinasse and molasses as mixed substrate. Process Biochem. 85, 12–18 (2019).

66. S. C. Reddington, M. Howarth, Secrets of a covalent interaction for biomaterials and biotechnology: SpyTag and SpyCatcher. Curr. Opin. Chem. Biol. 29, 94–99 (2015).

67. B. Zakeri, J. O. Fierer, E. Celik, E. C. Chittock, U. Schwarz-Linek, V. T. Moy, M. Howarth, Peptide tag forming a rapid covalent bond to a protein, through engineering a bacterial adhesin. Proc. Natl. Acad. Sci. U. S. A. 109, E690–697 (2012).

68. J. X. Wong, K. Ogura, S. Chen, B. H. A. Rehm, Bioengineered Polyhydroxyalkanoates as Immobilized Enzyme Scaffolds for Industrial Applications. Front. Bioeng. Biotechnol. 8, 156 (2020).

69. A. C. Jahns, B. H. A. Rehm, Tolerance of the Ralstonia eutropha Class I Polyhydroxyalkanoate Synthase for Translational Fusions to Its C Terminus Reveals a New Mode of Functional Display. Appl. Environ. Microbiol. 75, 5461–5466 (2009).

70. A. H. Keeble, P. Turkki, S. Stokes, I. N. A. Khairil Anuar, R. Rahikainen, V. P. Hytönen, M. Howarth, Approaching infinite affinity through engineering of peptide–protein interaction. Proc. Natl. Acad. Sci. 116, 26523–26533 (2019).

71. R. Karan, D. Renn, S. Nozue, L. Zhao, S. Habuchi, T. Allers, M. Rueping, Bioengineering of air-filled protein nanoparticles by genetic and chemical functionalization. J. Nanobiotechnology 21, 108 (2023).

72. R. Fan, J. Hakanpää, K. Elfving, H. Taberman, M. B. Linder, A. S. Aranko, Biomolecular Click Reactions Using a Minimal pH-Activated Catcher/Tag Pair for Producing Native-Sized Spider-Silk Proteins**. Angew. Chem. 135, e202216371 (2023).

73. W. Huang, A. Wilks, A rapid seamless method for gene knockout in Pseudomonas aeruginosa. BMC Microbiol. 17, 199 (2017).

74. Flowing Sotware documentation. https://bioscience.fi/cytometry-core/flowing-software/.

75. B. A. Nanes, Slide Set: Reproducible image analysis and batch processing with ImageJ. BioTechniques 59, 269–278 (2015).

76. H. Steel, R. Habgood, C. L. Kelly, A. Papachristodoulou, In situ characterisation and manipulation of biological systems with Chi.Bio. PLOS Biol. 18, e3000794 (2020).

