## Supplemental Material for "A Toolbox for Biomanufacturing of Functionalised PHA Nanoparticles with *C. necator*"

#### S1 Spot microdilutions in 96-well plate format

1. Cultures are diluted 1:10 along each of 8 plate columns.

2. A multi-channel pipette is then used to spot 5 µL from each column on an agar plate. The spots are left to dry completely before inverting the plates and placing them in the incubator.

3. Transformation efficiency can then be accurately quantified by counting the cells of the first countable spot on the agar plate.

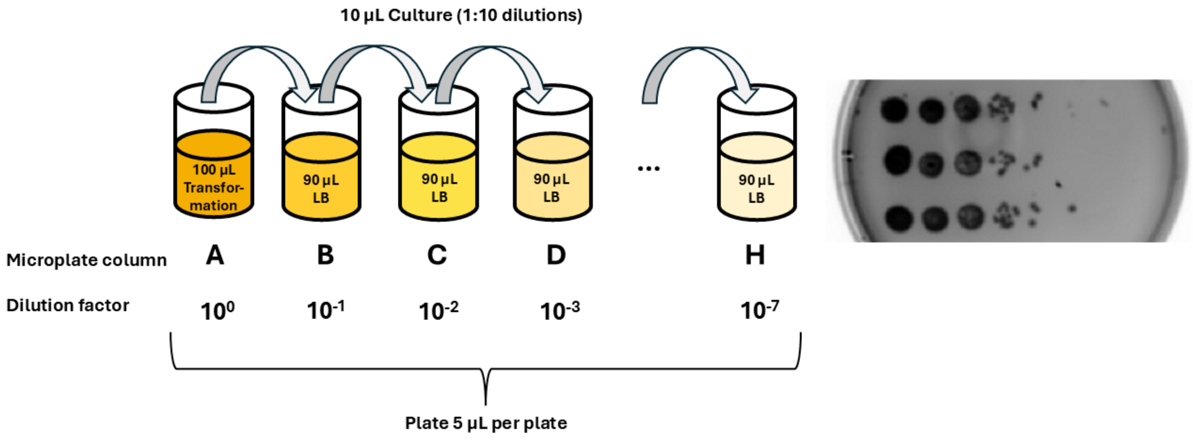


***Figure S1 – Spot microdilution procedure for quantifying transformation efficiency.***

#### S2 Primers used in Cloning

*Table S2.1 – Primer pairs for cloning phaC library and SpyCatch::SpyTag system.*

| PRIMER | PRIMER NAME | SEQUENCE | CONC |
|---|---|---|---|
| **PRH412_FWD** | pRH412_fwd | attatctagaggatccccgg | 25nm |
| **PRH412_REV** | pRH412_rev | ggttgtcctcctttgggg | 25nm |
| **PHAC_AC_WT_FWD** | AcPhaC_fwd | ccccccaaaggaggacaaccatgagccaatcatcttacgg | 25nm |
| **PHAC_AC_WT_REV** | AcPhaC_rev | ccggggatcctctagataattcatgcgggttcctcttc | 25nm |
| **PHAC_BS_WT_FWD** | BrPhaC_fwd | ccccccaaaggaggacaaccatggccaagactccccgaac | 25nm |
| **PHAC_BS_WT_REV** | BrPhaC_rev | ccggggatcctctagataattcaagacttcactttcacatagctgc | 25nm |
| **PHAC_CN_WT_FWD** | RePhaC_fwd | ccccccaaaggaggacaaccatggcgaccggcaaaggc | 25nm |
| **PHAC_CN_WT_REV** | RePhaC_rev | ccggggatcctctagataattcatgccttggctttgacgtatcg | 25nm |
| **PHAC_AC_M_FWD** | AcPhaC_evol_fwd | ccccccaaaggaggacaaccatgagccaatcatcttacgg | 25nm |
| **PHAC_AC_M_REV** | AcPhaC_evol_rev | ccggggatcctctagataattcatgcgggttcctcttc | 25nm |
| **PHAC_CN_M_FWD** | RePhac_F420S_fwd | ccccccaaaggaggacaaccatggcgaccggcaaaggc | 25nm |
| **PHAC_CN_M_REV** | RePhac_F420S_rev | ccggggatcctctagataattcatgccttggctttgacgtatcg | 25nm |

S3 Plasmids for phaC library and SpyCatch::SpyTag

Plasmid Sequences PhaC library

*pPhaC_Ac_wt*

### 1 attatctaga ggatccccgg gtaccgagct cgaattcgcg cggccgcggc ctaggcggcc

### 61 tcctgtgtga aattgttatc cgctttaatt aaaggcatca aataaaacga aaggctcagt

### 121 cgaaagactg ggcctttcgt tttatctgtt gtttgtcggt gaacgctctc ctgagtagga

### 181 caaatccgcc gccctagaca gctgggcgcg cccccctac gggcttgctc tccgggcttc

### 241 gccctgcgcg gtcgctgcgc tcccttgcca gcccgtggat atgtggacga tggccgcgag

### 301 cggccaccgg ctggctcgct tcgctcggcc cgtggacaac cctgctggac aagctgatgg

### 361 acaggctgcg cctgcccacg agcttgacca cagggattgc ccaccggcta cccagccttc

### 421 gaccacatac ccaccggctc caactgcgcg gcctgcggcc ttgccccatc aattttttta

### 481 attttctctg gggaaaagcc tccggcctgc ggcctgcgcg cttcgcttgc cggttggaca

### 541 ccaagtggaa ggcgggtcaa ggctcgcgca gcgaccgcgc agcggcttgg ccttgacgcg

### 601 cctggaacga cccaagccta tgcgagtggg ggcagtcgaa gggcgaagcc cgcccgcctg

### 661 ccccccgagc ctcacggcgg cgagtgcggg ggttccaagg gggcagcgcc accttgggca

### 721 aggccgaagg ccgcgcagtc gatcaacaag ccccggaggg gccacttttt gccggagggg

### 781 gagccgcgcc gaaggcgtgg gggaaccccg caggggtgcc cttctttggg caccaaagaa

### 841 ctagatatag ggcgaaatgc gaaagactta aaaatcaaca acttaaaaaa ggggggtacg

### 901 caacagctca ttgcggcacc ccccgcaata gctcattgcg taggttaaag aaaatctgta

### 961 attgactgcc acttttacgc aacgcataat tgttgtcgcg ctgccgaaaa gttgcagctg

### 1021 attgcgcatg gtgccgcaac cgtgcggcac ccctaccgca tggagataag catggccacg

### 1081 cagtccagag aaatcggcat tcaagccaag aacaagcccg gtcactgggt gcaaacggaa

### 1141 cgcaaagcgc atgaggcgtg ggccgggctt attgcgagga aacccacggc ggcaatgctg

### 1201 ctgcatcacc tcgtggcgca gatgggccac cagaacgccg tggtggtcag ccagaagaca

### 1261 ctttccaagc tcatcggacg ttctttgcgg acggtccaat acgcagtcaa ggacttggtg

### 1321 gccgagcgct ggatctccgt cgtgaagctc aacggccccg gcaccgtgtc ggcctacgtg

### 1381 gtcaatgacc gcgtggcgtg gggccagccc cgcgaccagt tgcgcctgtc ggtgttcagt

### 1441 gccgccgtgg tggttgatca cgacgaccag gacgaatcgc tgttggggca tggcgacctg

### 1501 cgccgcatcc cgaccctgta tccgggcgag cagcaactac cgaccggccc cggcgaggag

### 1561 ccgcccagcc agcccggcat tccgggcatg gaaccagacc tgccagcctt gaccgaaacg

### 1621 gaggaatggg aacggcgcgg gcagcagcgc ctgccgatgc ccgatgagcc gtgttttctg

### 1681 gacgatggcg agccgttgga gccgccgaca cgggtaacgc tgccgcgccg gtagggccgg

### 1741 cctacggcca gcctcgcaga gcaggattcc cgttgagcac cgccaggtgc gaataaggga

### 1801 cagtgaagaa ggaacacccg ctcgcgggtg ggcctacttc acctatcctg cccggctgac

### 1861 gccgttggat acaccaagga aagtctacac gaaccctttg gcaaaatcct gtatatcgtg

### 1921 cgaaaaagga tggatatacc gaaaaaatcg ctataatgac cccgaagcag ggttatgcag

### 1981 cggaaaagga caacgcgcgg accgcggtcc aattaattat tagaaaaatt catccagcat

### 2041 cagatgaaat tgcagtttgt tcatatccgg attatcaatg ccatatttct gaaacagacg

### 2101 tttttgcagg ctcgggctaa attcgcccag gcagttccac agaatggcca gatcctgata

### 2161 acgatccgca atgcccacac ggcccacatc aatgcagcca atcagtttgc cttcatcgaa

### 2221 aatcaggtta tccaggctaa aatcgccgtg ggtcaccacg ctatccgggc taaacggcag

### 2281 cagtttatgc atttctttcc acacctgttc caccggccag ccgttacgtt catcatcaaa

### 2341 atcgctcgca tccaccaggc cgttgttcat acggctctgc gcctgggcca gacgaaacac

### 2401 acgatcgctg ttaaacgggc agttgcacac cggaatgcta tgcagacgac gcagaaacac

### 2461 ggccagcgca tccacaatgt tttcgccgct atccggatat tcttccagca cctgaaacgc

### 2521 ggttttgccc ggaatcgcgg tggtcagcag ccacgcatca tccggggtgc gaataaaatg

### 2581 tttaatggtc ggcagcggca taaattcggt cagccagttc agacgcacca tttcatcggt

### 2641 cacatcgttc gccacgctgc ctttgccatg tttcagaaac agttccggcg catccggttt

### 2701 gccatacaga cgataaatgg tcgcgccgct ctgacccacg ttatcacgcg cccatttata

### 2761 gccatacaga tccgcatcca tgttgctgtt cagacgcgga cggctacagc tcgtttcacg

### 2821 ctgaatatgg ctcataacac cccttgtatt actgtttatg taagcagaca gttttattgt

### 2881 tcatgatgat atatttttat cttgtgcaat gtaacatcag agattttgag acacaaattt

### 2941 aaatcgtaat tattggggac ccctggattc tcaccaataa aaaacgcccg gcggcaaccg

### 3001 agcgttctga acaaatccag atggagttct gaggtcatta ctggatctat caacaggagt

### 3061 ccaagactag tcgccagggt tttcccagtc acgacgcggc cgcaagcttg catgcctgca

### 3121 ggttatgaca acttgacggc tacatcattc actttttctt cacaaccggc acggaactcg

### 3181 ctcgggctgg ccccggtgca ttttttaaat acccgcgaga aatagagttg atcgtcaaaa

### 3241 ccaacattgc gaccgacggt ggcgataggc atccgggtgg tgctcaaaag cagcttcgcc

### 3301 tggctgatac gttggtcctc gcgccagctt aagacgctaa tccctaactg ctggcggaaa

### 3361 agatgtgaca gacgcgacgg cgacaagcaa acatgctgtg cgacgctggc gatatcaaaa

### 3421 ttgctgtctg ccaggtgatc gctgatgtac tgacaagcct cgcgtacccg attatccatc

### 3481 ggtggatgga gcgactcgtt aatcgcttcc atgcgccgca gtaacaattg ctcaagcaga

### 3541 tttatcgcca gcagctccga atagcgccct tccccttgcc cggcgttaat gatttgccca

### 3601 aacaggtcgc tgaaatgcgg ctggtgcgct tcatccgggc gaaagaaccc cgtattggca

### 3661 aatattgacg gccagttaag ccattcatgc cagtaggcgc gcggacgaaa gtaaacccac

### 3721 tggtgatacc attcgcgagc ctccggatga cgaccgtagt gatgaatctc tcctggcggg

### 3781 aacagcaaaa tatcacccgg tcggcaaaca aattctcgtc cctgattttt caccaccccc

### 3841 tgaccgcgaa tggtgagatt gagaatataa cctttcattc ccagcggtcg gtcgataaaa

### 3901 aaatcgagat aaccgttggc ctcaatcggc gttaaacccg ccaccagatg ggcattaaac

### 3961 gagtatcccg gcagcagggg atcattttgc gcttcagcca tacttttcat actcccgcca

### 4021 ttcagagaag aaaccaattg tccatattgc atcagacatt gccgtcactg cgtcttttac

### 4081 tggctcttct cgctaaccaa accggtaacc ccgcttatta aaagcattct gtaacaaagc

### 4141 gggaccaaag ccatgacaaa aacgcgtaac aaaagtgtct ataatcacgg cagaaaagtc

### 4201 cacattgatt atttgcacgg cgtcacactt tgctatgcca tagcattttt atccataaga

### 4261 ttagcggatc ctacctgacg ctttttatcg caactctcta ctgtttctcc atgggccccc

### 4321 ccaaaggagg acaaccatga gccaatcatc ttacggcccg ctgttcgagg ccctggccca

### 4381 ctacaacgac aagctgctgg ccatggccag agcccagacc gagcgcacgg cccaggccct

### 4441 gctgcagacc agtctggacg atctgggcca ggtgctggag cagggcagcc agcagccctg

### 4501 gcagctgatc cagtcccaga tgaactggtg gcaggatcag ctcaagctga tgcagcacac

### 4561 cctgctcaag agtgcaggcc agccgagcga gccggtgatc accccggatc gcagcgatcg

### 4621 ccgcttcaag gccgaggcct ggagcgagca acccctctat gactacctca agcaatccta

### 4681 tctgctgacc gcaagacacc tgctggcggc ggtggatgcc ctggagggcg tgcccgagaa

### 4741 gagccgggag cgactgcgct ttttcacccg ccagtacgtc aacgccatgg ctcccagcaa

### 4801 cttcctggcc accaacccgg agctgctcaa gctcaccctg gagtccgacg gccagaatct

### 4861 ggtgcgcggc ctcgcgctcc tggccgaaga tctggagcgc agcgccgatc acctcaacat

### 4921 ccgcctgacc gacgagtccg ccttcgagct cggccgggat ctggcgacca ccccgggccg

### 4981 ggtagtgctg cgcaccgagc tctttgagct catccagtac cgcccgacca cggcgacggt

### 5041 gggcaagacc ccggtgctgg tcgtgccccc cttcatcaac aagtactaca tcatggacat

### 5101 gcgcccccag aactccctgg tggcctggct ggtcgcccag gggcagacgg tcttcatgat

### 5161 ctcctggcgc aacccggggg tggcacaggg gctggtcgat ctcgacgact acgtggtgga

### 5221 cggggtcatc gccgccctgg acgccgtgga ggcggccacc ggcgagcggg aggtgcacgg

### 5281 catcggctac tgcatcggtg gcaccgcatt gtcactcgcc atgggctggc tggcagcgcg

### 5341 gcgccagaag cagcgggtgc gtagcgccac cctgttcacc accctgctgg acttctccca

### 5401 gcccggggag cttggcatct tcatccacga gcccatcatc gcggcgctcg aggcgcagaa

### 5461 cgaggccagg ggcatcatgg acgggcgcca gctggcggtc tccttcagtc tgctgcggga

### 5521 gaacagcctc tactggaact actacatcga cagctacctc aagggtcaga gcccggtggc

### 5581 gttcgatctg ctgcactgga acagcgacag caccaacgtg gcgggcaaga cccacaacag

### 5641 cctgctgcgc cgactctacc tggaaaacca gctggtgaag ggggagctca agatccgcca

### 5701 cacccgcatc gatctcgcca aggtgaagac cccggtgctg ctggtgtccg ccgtggacga

### 5761 tcacatcgcc ctctggcagg gaacctggca gggcatggcg ctgttcgggg gcgagcggcg

### 5821 cttcattctg gccgaatccg ggcacatcgc cggcatcatc aatccgccgg atgccaacaa

### 5881 atacggtttc tggcagagcg aggccgaggc ggacagcccg gcgcagtggc tggcgggggc

### 5941 gacccaccag agcggctcct ggtggcccga gatgatgcgc tttatcaaag accgtgacga

### 6001 ggcagagccc gtccccgccc agcagcctgc cgaggggttg gaggcggccc ccggcagcta

### 6061 cgtcaaggta cggctcaatc cggtgtttgc cggcatcagg aagcaagaag aggaacccgc

### 6121 atga

*pPHAC_Bs_wt*

### 1 attatctaga ggatccccgg gtaccgagct cgaattcgcg cggccgcggc ctaggcggcc

### 61 tcctgtgtga aattgttatc cgctttaatt aaaggcatca aataaaacga aaggctcagt

### 121 cgaaagactg ggcctttcgt tttatctgtt gtttgtcggt gaacgctctc ctgagtagga

### 181 caaatccgcc gccctagaca gctgggcgcg cccccctac gggcttgctc tccgggcttc

### 241 gccctgcgcg gtcgctgcgc tcccttgcca gcccgtggat atgtggacga tggccgcgag

### 301 cggccaccgg ctggctcgct tcgctcggcc cgtggacaac cctgctggac aagctgatgg

### 361 acaggctgcg cctgcccacg agcttgacca cagggattgc ccaccggcta cccagccttc

### 421 gaccacatac ccaccggctc caactgcgcg gcctgcggcc ttgccccatc aattttttta

### 481 attttctctg gggaaaagcc tccggcctgc ggcctgcgcg cttcgcttgc cggttggaca

### 541 ccaagtggaa ggcgggtcaa ggctcgcgca gcgaccgcgc agcggcttgg ccttgacgcg

### 601 cctggaacga cccaagccta tgcgagtggg ggcagtcgaa gggcgaagcc cgcccgcctg

### 661 ccccccgagc ctcacggcgg cgagtgcggg ggttccaagg gggcagcgcc accttgggca

### 721 aggccgaagg ccgcgcagtc gatcaacaag ccccggaggg gccacttttt gccggagggg

### 781 gagccgcgcc gaaggcgtgg gggaaccccg caggggtgcc cttctttggg caccaaagaa

### 841 ctagatatag ggcgaaatgc gaaagactta aaaatcaaca acttaaaaaa ggggggtacg

### 901 caacagctca ttgcggcacc ccccgcaata gctcattgcg taggttaaag aaaatctgta

### 961 attgactgcc acttttacgc aacgcataat tgttgtcgcg ctgccgaaaa gttgcagctg

### 1021 attgcgcatg gtgccgcaac cgtgcggcac ccctaccgca tggagataag catggccacg

### 1081 cagtccagag aaatcggcat tcaagccaag aacaagcccg gtcactgggt gcaaacggaa

### 1141 cgcaaagcgc atgaggcgtg ggccgggctt attgcgagga aacccacggc ggcaatgctg

### 1201 ctgcatcacc tcgtggcgca gatgggccac cagaacgccg tggtggtcag ccagaagaca

### 1261 ctttccaagc tcatcggacg ttctttgcgg acggtccaat acgcagtcaa ggacttggtg

### 1321 gccgagcgct ggatctccgt cgtgaagctc aacggccccg gcaccgtgtc ggcctacgtg

### 1381 gtcaatgacc gcgtggcgtg gggccagccc cgcgaccagt tgcgcctgtc ggtgttcagt

### 1441 gccgccgtgg tggttgatca cgacgaccag gacgaatcgc tgttggggca tggcgacctg

### 1501 cgccgcatcc cgaccctgta tccgggcgag cagcaactac cgaccggccc cggcgaggag

### 1561 ccgcccagcc agcccggcat tccgggcatg gaaccagacc tgccagcctt gaccgaaacg

### 1621 gaggaatggg aacggcgcgg gcagcagcgc ctgccgatgc ccgatgagcc gtgttttctg

### 1681 gacgatggcg agccgttgga gccgccgaca cgggtaacgc tgccgcgccg gtagggccgg

### 1741 cctacggcca gcctcgcaga gcaggattcc cgttgagcac cgccaggtgc gaataaggga

### 1801 cagtgaagaa ggaacacccg ctcgcgggtg ggcctacttc acctatcctg cccggctgac

### 1861 gccgttggat acaccaagga aagtctacac gaaccctttg gcaaaatcct gtatatcgtg

### 1921 cgaaaaagga tggatatacc gaaaaaatcg ctataatgac cccgaagcag ggttatgcag

### 1981 cggaaaagga caacgcgcgg accgcggtcc aattaattat tagaaaaatt catccagcat

### 2041 cagatgaaat tgcagtttgt tcatatccgg attatcaatg ccatatttct gaaacagacg

### 2101 tttttgcagg ctcgggctaa attcgcccag gcagttccac agaatggcca gatcctgata

### 2161 acgatccgca atgcccacac ggcccacatc aatgcagcca atcagtttgc cttcatcgaa

### 2221 aatcaggtta tccaggctaa aatcgccgtg ggtcaccacg ctatccgggc taaacggcag

### 2281 cagtttatgc atttctttcc acacctgttc caccggccag ccgttacgtt catcatcaaa

### 2341 atcgctcgca tccaccaggc cgttgttcat acggctctgc gcctgggcca gacgaaacac

### 2401 acgatcgctg ttaaacgggc agttgcacac cggaatgcta tgcagacgac gcagaaacac

### 2461 ggccagcgca tccacaatgt tttcgccgct atccggatat tcttccagca cctgaaacgc

### 2521 ggttttgccc ggaatcgcgg tggtcagcag ccacgcatca tccggggtgc gaataaaatg

### 2581 tttaatggtc ggcagcggca taaattcggt cagccagttc agacgcacca tttcatcggt

### 2641 cacatcgttc gccacgctgc ctttgccatg tttcagaaac agttccggcg catccggttt

### 2701 gccatacaga cgataaatgg tcgcgccgct ctgacccacg ttatcacgcg cccatttata

### 2761 gccatacaga tccgcatcca tgttgctgtt cagacgcgga cggctacagc tcgtttcacg

### 2821 ctgaatatgg ctcataacac cccttgtatt actgtttatg taagcagaca gttttattgt

### 2881 tcatgatgat atatttttat cttgtgcaat gtaacatcag agattttgag acacaaattt

### 2941 aaatcgtaat tattggggac ccctggattc tcaccaataa aaaacgcccg gcggcaaccg

### 3001 agcgttctga acaaatccag atggagttct gaggtcatta ctggatctat caacaggagt

### 3061 ccaagactag tcgccagggt tttcccagtc acgacgcggc cgcaagcttg catgcctgca

### 3121 ggttatgaca acttgacggc tacatcattc actttttctt cacaaccggc acggaactcg

### 3181 ctcgggctgg ccccggtgca ttttttaaat acccgcgaga aatagagttg atcgtcaaaa

### 3241 ccaacattgc gaccgacggt ggcgataggc atccgggtgg tgctcaaaag cagcttcgcc

### 3301 tggctgatac gttggtcctc gcgccagctt aagacgctaa tccctaactg ctggcggaaa

### 3361 agatgtgaca gacgcgacgg cgacaagcaa acatgctgtg cgacgctggc gatatcaaaa

### 3421 ttgctgtctg ccaggtgatc gctgatgtac tgacaagcct cgcgtacccg attatccatc

### 3481 ggtggatgga gcgactcgtt aatcgcttcc atgcgccgca gtaacaattg ctcaagcaga

### 3541 tttatcgcca gcagctccga atagcgccct tccccttgcc cggcgttaat gatttgccca

### 3601 aacaggtcgc tgaaatgcgg ctggtgcgct tcatccgggc gaaagaaccc cgtattggca

### 3661 aatattgacg gccagttaag ccattcatgc cagtaggcgc gcggacgaaa gtaaacccac
### 3721 tggtgatacc attcgcgagc ctccggatga cgaccgtagt gatgaatctc tcctggcggg
### 3781 aacagcaaaa tatcacccgg tcggcaaaca aattctcgtc cctgattttt caccaccccc
### 3841 tgaccgcgaa tggtgagatt gagaatataa cctttcattc ccagcggtcg gtcgataaaa
### 3901 aaatcgagat aaccgttggc ctcaatcggc gttaaacccg ccaccagatg ggcattaaac
### 3961 gagtatcccg gcagcagggg atcattttgc gcttcagcca tacttttcat actcccgcca
### 4021 ttcagagaag aaaccaattg tccatattgc atcagacatt gccgtcactg cgtcttttac
### 4081 tggctcttct cgctaaccaa accggtaacc ccgcttatta aaagcattct gtaacaaagc
### 4141 gggaccaaag ccatgacaaa aacgcgtaac aaaagtgtct ataatcacgg cagaaaagtc
### 4201 cacattgatt atttgcacgg cgtcacactt tgctatgcca tagcattttt atccataaga
### 4261 ttagcggatc ctacctgacg ctttttatcg caactctcta ctgtttctcc atgggccccc
### 4321 ccaaaggagg acaaccatgg ccaagactcc ccgaacgccg acagacgccc ccgatcgccc
### 4381 cgcacgcaag gccggacgtc cgacgtccgc caagccccgc ccgcctcggc agcctaagtc
### 4441 gaaacctgcc gaaacgcggc aagccgcctc tgaaacccat gcagcgccga ctcctgaacc
### 4501 gaccgaagcg acacccaatc aggcccaatt gatcgaaacc ctgtcgatga atctggccaa
### 4561 ggcggcaatg atggcccaga gcgcaatcgc cgaggcggcg ctgacccagg ccgatcggcc
### 4621 ggccgccctt tccgccgatc ccttcaacgt cgcgccggcc atgacgtcgg tcatgaccag
### 4681 cctggccgcc cagcccgaca aaatgattca ggcccaggcc gacctgtttg gccgctatat
### 4741 gcagctctgg tcatcgacgg cgcgtcaggc ggccggcgag gcgcccgatc ccgcccccggt
### 4801 cgacaagcgg ttcaaggacc cggcttggtc cgaaaacccc atgttcgaca tgatgcggcg
### 4861 atcctatctg ctgacgtcgg actggatgaa cggtctgatc gccggggtcg aggacgtcga
### 4921 tccgacgctg aaacgtcgcg cccaattctt tacccgcctg ctgaccgacg ccttctcgcc
### 4981 gtccaacttt ttggcctcca accccgtcgc gctgaaggcc ttggcggaga ccagcggtga
### 5041 atcgctggtg aaaggtatgc agaacttcgc cgccgacttg gaacgtggcg gcggatcgct

### 5101 gcggatcagc caggccgact acggaaagtt cgtcgtcggc gagaacgtgg ccaccgcgcc

### 5161 gggccaggtc gtctggcgcg atgagctgtt cgaactgatc cagtacgcac cctcgactga

### 5221 gagtcagcac gacatcccgc tgctgatctt cccgccctgg atcaacaaat tctacatcat

### 5281 ggacctgcag ccggcgaact cgctgatccg ctggctgtcg gctcagggct tcacggtctt

### 5341 cgtctgttcc tgggtcaatc cggacaggga taaggccgga ttcggcttcg acgactatct

### 5401 ggacaagggc atctatcgcg cggtcgagaa gacgctggag caggctggca cgaagcaact

### 5461 gaacgccgtg ggctattgca tcggcggcac cctgcttggc gctggcctgg cgcacatggc

### 5521 ggccaagggc gacaagcgca tcgccgccgc taccttcttc gccgctcaac atgacttcgc

### 5581 cgaggccggc gacctgcttc tgttcaccga cgaacactgg atcgccgaga tcgagcgtca

### 5641 gatggatgcg gcgggcggcg tcctgccggg cgcggccatg gcggaaacgt tcaacgccct

### 5701 gcggtccaat gacctgatct ggtccttttt cattagcaac tatctgctgg ggaaggatcc

### 5761 gcccgccttc gacctgttgt tctggaacgc cgatcagacc cgcatgccca aggcgctgca

### 5821 tctggactat ctgcgtcaga tgtatggggc caacgccctg gccaaggggc agttcgagat

### 5881 cggcggcctg acggcggacc tgtcgaaggt cgagatcccg ctctattttc aggccagccg

### 5941 agaggatcac atcgcgccga tgaactcggt ctatcgatcc gccaaactgt tcggcggcaa

### 6001 ggatgtgacc ttcaccctgg ccgggtcagg tcacatcgcc ggcgtcatca acgcccccgc

### 6061 cgcaaagaag tatcagcatt ggaccaaccc tgccctgccc gcgaccttgg ccgaatggca

### 6121 agctgacgcc gtcgaacatc ccggcagctg gtgggagcat tggtcggcct ggctcggcgc

### 6181 gcgttccggc gctcaaatcc ccgcccgcaa ccccgccaag ggcccgctca agcccatcga

### 6241 gccggcgccg ggcagctatg tgaaagtgaa gtcttga

*pPhaC_Cn_wt*

1 attatctaga ggatccccgg gtaccgagct cgaattcgcg cggccgcggc ctaggcggcc

### 61 tcctgtgtga aattgttatc cgctttaatt aaaggcatca aataaaacga aaggctcagt

### 121 cgaaagactg ggcctttcgt tttatctgtt gtttgtcggt gaacgctctc ctgagtagga

### 181 caaatccgcc gccctagaca gctgggcgcg cccccctac gggcttgctc tccgggcttc

### 241 gccctgcgcg gtcgctgcgc tcccttgcca gcccgtggat atgtggacga tggccgcgag

### 301 cggccaccgg ctggctcgct tcgctcggcc cgtggacaac cctgctggac aagctgatgg

### 361 acaggctgcg cctgcccacg agcttgacca cagggattgc ccaccggcta cccagccttc

### 421 gaccacatac ccaccggctc caactgcgcg gcctgcggcc ttgccccatc aattttttta

### 481 attttctctg gggaaaagcc tccggcctgc ggcctgcgcg cttcgcttgc cggttggaca

### 541 ccaagtggaa ggcgggtcaa ggctcgcgca gcgaccgcgc agcggcttgg ccttgacgcg

### 601 cctggaacga cccaagccta tgcgagtggg ggcagtcgaa gggcgaagcc cgcccgcctg

### 661 ccccccgagc ctcacggcgg cgagtgcggg ggttccaagg gggcagcgcc accttgggca

### 721 aggccgaagg ccgcgcagtc gatcaacaag ccccggaggg gccacttttt gccggagggg

### 781 gagccgcgcc gaaggcgtgg gggaaccccg caggggtgcc cttctttggg caccaaagaa

### 841 ctagatatag ggcgaaatgc gaaagactta aaaatcaaca acttaaaaaa ggggggtacg

### 901 caacagctca ttgcggcacc ccccgcaata gctcattgcg taggttaaag aaaatctgta

### 961 attgactgcc acttttacgc aacgcataat tgttgtcgcg ctgccgaaaa gttgcagctg

### 1021 attgcgcatg gtgccgcaac cgtgcggcac ccctaccgca tggagataag catggccacg

### 1081 cagtccagag aaatcggcat tcaagccaag aacaagcccg gtcactgggt gcaaacggaa

### 1141 cgcaaagcgc atgaggcgtg ggccgggctt attgcgagga aacccacggc ggcaatgctg

### 1201 ctgcatcacc tcgtggcgca gatgggccac cagaacgccg tggtggtcag ccagaagaca

### 1261 ctttccaagc tcatcggacg ttctttgcgg acggtccaat acgcagtcaa ggacttggtg

### 1321 gccgagcgct ggatctccgt cgtgaagctc aacggccccg gcaccgtgtc ggcctacgtg

### 1381 gtcaatgacc gcgtggcgtg ggccagccc cgcgaccagt tgcgcctgtc ggtgttcagt

### 1441 gccgccgtgg tggttgatca cgacgaccag gacgaatcgc tgttggggca tggcgacctg

### 1501 cgccgcatcc cgaccctgta tccgggcgag cagcaactac cgaccggccc cggcgaggag

### 1561 ccgcccagcc agcccggcat tccgggcatg gaaccagacc tgccagcctt gaccgaaacg

### 1621 gaggaatggg aacggcgcgg gcagcagcgc ctgccgatgc ccgatgagcc gtgttttctg

### 1681 gacgatggcg agccgttgga gccgccgaca cgggtaacgc tgccgcgccg gtagggccgg

### 1741 cctacggcca gcctcgcaga gcaggattcc cgttgagcac cgccaggtgc gaataaggga

### 1801 cagtgaagaa ggaacacccg ctcgcgggtg ggcctacttc acctatcctg cccggctgac

### 1861 gccgttggat acaccaagga aagtctacac gaaccctttg gcaaaatcct gtatatcgtg

### 1921 cgaaaaagga tggatatacc gaaaaaatcg ctataatgac cccgaagcag ggttatgcag

### 1981 cggaaaagga caacgcgcgg accgcggtcc aattaattat tagaaaaatt catccagcat

### 2041 cagatgaaat tgcagtttgt tcatatccgg attatcaatg ccatatttct gaaacagacg

### 2101 tttttgcagg ctcgggctaa attcgcccag gcagttccac agaatggcca gatcctgata

### 2161 acgatccgca atgcccacac ggcccacatc aatgcagcca atcagtttgc cttcatcgaa

### 2221 aatcaggtta tccaggctaa aatcgccgtg ggtcaccacg ctatccgggc taaacggcag

### 2281 cagtttatgc atttctttcc acacctgttc caccggccag ccgttacgtt catcatcaaa

### 2341 atcgctcgca tccaccaggc cgttgttcat acggctctgc gcctgggcca gacgaaacac

### 2401 acgatcgctg ttaaacgggc agttgcacac cggaatgcta tgcagacgac gcagaaacac

### 2461 ggccagcgca tccacaatgt tttcgccgct atccggatat tcttccagca cctgaaacgc

### 2521 ggttttgccc ggaatcgcgg tggtcagcag ccacgcatca tccggggtgc gaataaaatg

### 2581 tttaatggtc ggcagcggca taaattcggt cagccagttc agacgcacca tttcatcggt

### 2641 cacatcgttc gccacgctgc ctttgccatg tttcagaaac agttccggcg catccggttt

### 2701 gccatacaga cgataaatgg tcgcgccgct ctgacccacg ttatcacgcg cccatttata

### 2761 gccatacaga tccgcatcca tgttgctgtt cagacgcgga cggctacagc tcgtttcacg

### 2821 ctgaatatgg ctcataacac cccttgtatt actgtttatg taagcagaca gttttattgt

### 2881 tcatgatgat atatttttat cttgtgcaat gtaacatcag agattttgag acacaaattt

### 2941 aaatcgtaat tattggggac ccctggattc tcaccaataa aaaacgcccg gcggcaaccg

### 3001 agcgttctga acaaatccag atggagttct gaggtcatta ctggatctat caacaggagt

### 3061 ccaagactag tcgccagggt tttcccagtc acgacgcggc cgcaagcttg catgcctgca

### 3121 ggttatgaca acttgacggc tacatcattc actttttctt cacaaccggc acggaactcg

### 3181 ctcgggctgg ccccggtgca ttttttaaat acccgcgaga aatagagttg atcgtcaaaa

### 3241 ccaacattgc gaccgacggt ggcgataggc atccgggtgg tgctcaaaag cagcttcgcc

### 3301 tggctgatac gttggtcctc gcgccagctt aagacgctaa tccctaactg ctggcggaaa

### 3361 agatgtgaca gacgcgacgg cgacaagcaa acatgctgtg cgacgctggc gatatcaaaa

### 3421 ttgctgtctg ccaggtgatc gctgatgtac tgacaagcct cgcgtacccg attatccatc

### 3481 ggtggatgga gcgactcgtt aatcgcttcc atgcgccgca gtaacaattg ctcaagcaga

### 3541 tttatcgcca gcagctccga atagcgccct tccccttgcc cggcgttaat gatttgccca

### 3601 aacaggtcgc tgaaatgcgg ctggtgcgct tcatccgggc gaaagaaccc cgtattggca

### 3661 aatattgacg gccagttaag ccattcatgc cagtaggcgc gcggacgaaa gtaaacccac

### 3721 tggtgatacc attcgcgagc ctccggatga cgaccgtagt gatgaatctc tcctggcggg

### 3781 aacagcaaaa tatcacccgg tcggcaaaca aattctcgtc cctgattttt caccaccccc

### 3841 tgaccgcgaa tggtgagatt gagaatataa cctttcattc ccagcggtcg gtcgataaaa

### 3901 aaatcgagat aaccgttggc ctcaatcggc gttaaacccg ccaccagatg ggcattaaac

### 3961 gagtatcccg gcagcagggg atcattttgc gcttcagcca tacttttcat actcccgcca

### 4021 ttcagagaag aaaccaattg tccatattgc atcagacatt gccgtcactg cgtcttttac

### 4081 tggctcttct cgctaaccaa accggtaacc ccgcttatta aaagcattct gtaacaaagc

### 4141 gggaccaaag ccatgacaaa aacgcgtaac aaaagtgtct ataatcacgg cagaaaagtc

### 4201 cacattgatt atttgcacgg cgtcacactt tgctatgcca tagcattttt atccataaga

### 4261 ttagcggatc ctacctgacg ctttttatcg caactctcta ctgtttctcc atgggccccc

### 4321 ccaaaggagg acaaccatgg cgaccggcaa aggcgcggca gcttccacgc aggaaggcaa

### 4381 gtcccaacca ttcaaggtca cgccggggcc attcgatcca gccacatggc tggaatggtc

### 4441 ccgccagtgg cagggcactg aaggcaacgg ccacgcggcc gcgtccggca ttccgggcct

### 4501 ggatgcgctg gcaggcgtca agatcgcgcc ggcgcagctg ggtgatatcc agcagcgcta

### 4561 catgaaggac ttctcagcgc tgtggcaggc catggccgag ggcaaggccg aggccaccgg

### 4621 tccgctgcac gaccggcgct tcgccggcga cgcatggcgc accaacctcc catatcgctt

### 4681 cgctgccgcg ttctacctgc tcaatgcgcg cgccttgacc gagctggccg atgccgtcga

### 4741 ggccgatgcc aagacccgcc agcgcatccg cttcgcgatc tcgcaatggg tcgatgcgat

### 4801 gtcgcccgcc aacttccttg ccaccaatcc cgaggcgcag cgcctgctga tcgagtcggg

### 4861 cggcgaatcg ctgcgtgccg gcgtgcgcaa catgatggaa gacctgacac gcggcaagat

### 4921 ctcgcagacc gacgagagcg cgtttgaggt cggccgcaat gtcgcggtga ccgaaggcgc

### 4981 cgtggtcttc gagaacgagt acttccagct gttgcagtac aagccgctga ccgacaaggt

### 5041 gcacgcgcgc ccgctgctga tggtgccgcc gtgcatcaac aagtactaca tcctggacct

### 5101 gcagccggag agctcgctgg tgcgccatgt ggtggagcag ggacatacgg tgtttctggt

### 5161 gtcgtggcgc aatccggacg ccagcatggc cggcagcacc tgggacgact acatcgagca

### 5221 cgcggccatc cgcgccatcg aagtcgcgcg cgacatcagc ggccaggaca agatcaacgt

### 5281 gctcggcttc tgcgtgggcg gcaccattgt ctcgaccgcg ctggcggtgc tggccgcgcg

### 5341 cggcgagcac ccggccgcca gcgtcacgct gctgaccacg ctgctggact ttgccgacac

### 5401 gggcatcctc gacgtctttg tcgacgaggg ccatgtgcag ttgcgcgagg ccacgctggg

### 5461 cggcggcgcc ggcgcgccgt gcgcgctgct gcgcggcctt gagctggcca ataccttctc

### 5521 gttcttgcgc ccgaacgacc tggtgtggaa ctacgtggtc gacaactacc tgaagggcaa

### 5581 cacgccggtg ccgttcgacc tgctgttctg gaacggcgac gccaccaacc tgccggggcc

### 5641 gtggtactgc tggtacctgc gccacaccta cctgcagaac gagctcaagg taccgggcaa

### 5701 gctgaccgtg tgcggcgtgc cggtggacct ggccagcatc gacgtgccga cctatatcta

### 5761 cggctcgcgc gaagaccata tcgtgccgtg gaccgcggcc tatgcctcga ccgcgctgct

### 5821 ggcgaacaag ctgcgcttcg tgctgggtgc gtcgggccat atcgccggtg tgatcaaccc

### 5881 gccggccaag aacaagcgca gccactggac taacgatgcg ctgccggagt cgccgcagca

### 5941 atggctggcc ggcgccatcg agcatcacgg cagctggtgg ccggactgga ccgcatggct

### 6001 ggccgggcag gccggcgcga aacgcgccgc gcccgccaac tatggcaatg cgcgctatcg

### 6061 cgcaatcgaa cccgcgcctg ggcgatacgt caaagccaag gcatga

*pPHA_Cn_m*

### 1 attatctaga ggatccccgg gtaccgagct cgaattcgcg cggccgcggc ctaggcggcc

### 61 tcctgtgtga aattgttatc cgctttaatt aaaggcatca aataaaacga aaggctcagt

### 121 cgaaagactg ggcctttcgt tttatctgtt gtttgtcggt gaacgctctc ctgagtagga

### 181 caaatccgcc gccctagaca gctgggcgcg cccccctac gggcttgctc tccgggcttc

### 241 gccctgcgcg gtcgctgcgc tcccttgcca gcccgtggat atgtggacga tggccgcgag

### 301 cggccaccgg ctggctcgct tcgctcggcc cgtggacaac cctgctggac aagctgatgg

### 361 acaggctgcg cctgcccacg agcttgacca cagggattgc ccaccggcta cccagccttc

### 421 gaccacatac ccaccggctc caactgcgcg gcctgcggcc ttgccccatc aatttttta

### 481 attttctctg gggaaaagcc tccggcctgc ggcctgcgcg cttcgcttgc cggttggaca

### 541 ccaagtggaa ggcgggtcaa ggctcgcgca gcgaccgcgc agcggcttgg ccttgacgcg

### 601 cctggaacga cccaagccta tgcgagtggg ggcagtcgaa gggcgaagcc cgcccgcctg

### 661 ccccccgagc ctcacggcgg cgagtgcggg ggttccaagg gggcagcgcc accttgggca

### 721 aggccgaagg ccgcgcagtc gatcaacaag ccccggaggg gccacttttt gccggagggg

### 781 gagccgcgcc gaaggcgtgg gggaaccccg caggggtgcc cttctttggg caccaaagaa

### 841 ctagatatag ggcgaaatgc gaaagactta aaaatcaaca acttaaaaaa gggggtacg

### 901 caacagctca ttgcggcacc ccccgcaata gctcattgcg taggttaaag aaaatctgta

### 961 attgactgcc acttttacgc aacgcataat tgttgtcgcg ctgccgaaaa gttgcagctg

### 1021 attgcgcatg gtgccgcaac cgtgcggcac ccctaccgca tggagataag catggccacg

### 1081 cagtccagag aaatcggcat tcaagccaag aacaagcccg gtcactgggt gcaaacggaa

### 1141 cgcaaagcgc atgaggcgtg ggccgggctt attgcgagga aacccacggc ggcaatgctg

### 1201 ctgcatcacc tcgtggcgca gatgggccac cagaacgccg tggtggtcag ccagaagaca

### 1261 ctttccaagc tcatcggacg ttctttgcgg acggtccaat acgcagtcaa ggacttggtg

### 1321 gccgagcgct ggatctccgt cgtgaagctc aacggccccg gcaccgtgtc ggcctacgtg

### 1381 gtcaatgacc gcgtggcgtg ggccagccc cgcgaccagt tgcgcctgtc ggtgttcagt

### 1441 gccgccgtgg tggttgatca cgacgaccag gacgaatcgc tgttggggca tggcgacctg

### 1501 cgccgcatcc cgaccctgta tccgggcgag cagcaactac cgaccggccc cggcgaggag

### 1561 ccgcccagcc agcccggcat tccgggcatg gaaccagacc tgccagcctt gaccgaaacg

### 1621 gaggaatggg aacggcgcgg gcagcagcgc ctgccgatgc ccgatgagcc gtgttttctg

### 1681 gacgatggcg agccgttgga gccgccgaca cgggtaacgc tgccgcgccg gtagggccgg

### 1741 cctacggcca gcctcgcaga gcaggattcc cgttgagcac cgccaggtgc gaataaggga

### 1801 cagtgaagaa ggaacacccg ctcgcgggtg ggcctacttc acctatcctg cccggctgac

### 1861 gccgttggat acaccaagga aagtctacac gaaccctttg gcaaaatcct gtatatcgtg

### 1921 cgaaaaagga tggatatacc gaaaaaatcg ctataatgac cccgaagcag ggttatgcag

### 1981 cggaaaagga caacgcgcgg accgcggtcc aattaattat tagaaaaatt catccagcat

### 2041 cagatgaaat tgcagtttgt tcatatccgg attatcaatg ccatatttct gaaacagacg

### 2101 tttttgcagg ctcgggctaa attcgcccag gcagttccac agaatggcca gatcctgata

### 2161 acgatccgca atgcccacac ggcccacatc aatgcagcca atcagtttgc cttcatcgaa

### 2221 aatcaggtta tccaggctaa aatcgccgtg ggtcaccacg ctatccgggc taaacggcag

### 2281 cagtttatgc atttctttcc acacctgttc caccggccag ccgttacgtt catcatcaaa

### 2341 atcgctcgca tccaccaggc cgttgttcat acggctctgc gcctgggcca gacgaaacac

### 2401 acgatcgctg ttaaacgggc agttgcacac cggaatgcta tgcagacgac gcagaaacac

### 2461 ggccagcgca tccacaatgt tttcgccgct atccggatat tcttccagca cctgaaacgc

### 2521 ggttttgccc ggaatcgcgg tggtcagcag ccacgcatca tccggggtgc gaataaaatg

### 2581 tttaatggtc ggcagcggca taaattcggt cagccagttc agacgcacca tttcatcggt

### 2641 cacatcgttc gccacgctgc ctttgccatg tttcagaaac agttccggcg catccggttt

### 2701 gccatacaga cgataaatgg tcgcgccgct ctgacccacg ttatcacgcg cccatttata

### 2761 gccatacaga tccgcatcca tgttgctgtt cagacgcgga cggctacagc tcgtttcacg

### 2821 ctgaatatgg ctcataacac cccttgtatt actgtttatg taagcagaca gttttattgt

### 2881 tcatgatgat atatttttat cttgtgcaat gtaacatcag agattttgag acacaaattt

### 2941 aaatcgtaat tattggggac ccctggattc tcaccaataa aaaacgcccg gcggcaaccg

### 3001 agcgttctga acaaatccag atggagttct gaggtcatta ctggatctat caacaggagt

### 3061 ccaagactag tcgccagggt tttcccagtc acgacgcggc cgcaagcttg catgcctgca

### 3121 ggttatgaca acttgacggc tacatcattc actttttctt cacaaccggc acggaactcg

### 3181 ctcgggctgg ccccggtgca ttttttaaat acccgcgaga aatagagttg atcgtcaaaa

### 3241 ccaacattgc gaccgacggt ggcgataggc atccgggtgg tgctcaaaag cagcttcgcc

### 3301 tggctgatac gttggtcctc gcgccagctt aagacgctaa tccctaactg ctggcggaaa

### 3361 agatgtgaca gacgcgacgg cgacaagcaa acatgctgtg cgacgctggc gatatcaaaa

### 3421 ttgctgtctg ccaggtgatc gctgatgtac tgacaagcct cgcgtacccg attatccatc

### 3481 ggtggatgga gcgactcgtt aatcgcttcc atgcgccgca gtaacaattg ctcaagcaga

### 3541 tttatcgcca gcagctccga atagcgccct tccccttgcc cggcgttaat gatttgccca

### 3601 aacaggtcgc tgaaatgcgg ctggtgcgct tcatccgggc gaaagaaccc cgtattggca

### 3661 aatattgacg gccagttaag ccattcatgc cagtaggcgc gcggacgaaa gtaaacccac

### 3721 tggtgatacc attcgcgagc ctccggatga cgaccgtagt gatgaatctc tcctggcggg

### 3781 aacagcaaaa tatcacccgg tcggcaaaca aattctcgtc cctgattttt caccaccccc

### 3841 tgaccgcgaa tggtgagatt gagaatataa cctttcattc ccagcggtcg gtcgataaaa

### 3901 aaatcgagat aaccgttggc ctcaatcggc gttaaacccg ccaccagatg ggcattaaac

### 3961 gagtatcccg gcagcagggg atcattttgc gcttcagcca tacttttcat actcccgcca

### 4021 ttcagagaag aaaccaattg tccatattgc atcagacatt gccgtcactg cgtcttttac

### 4081 tggctcttct cgctaaccaa accggtaacc ccgcttatta aaagcattct gtaacaaagc

### 4141 gggaccaaag ccatgacaaa aacgcgtaac aaaagtgtct ataatcacgg cagaaaagtc

### 4201 cacattgatt atttgcacgg cgtcacactt tgctatgcca tagcattttt atccataaga

### 4261 ttagcggatc ctacctgacg ctttttatcg caactctcta ctgtttctcc atgggccccc

### 4321 ccaaaggagg acaaccatgg cgaccggcaa aggcgcggca gcttccacgc aggaaggcaa

### 4381 gtcccaacca ttcaaggtca cgccggggcc attcgatcca gccacatggc tggaatggtc

### 4441 ccgccagtgg cagggcactg aaggcaacgg ccacgcggcc gcgtccggca ttccgggcct

### 4501 ggatgcgctg gcaggcgtca agatcgcgcc ggcgcagctg ggtgatatcc agcagcgcta

### 4561 catgaaggac ttctcagcgc tgtggcaggc catggccgag ggcaaggccg aggccaccgg

### 4621 tccgctgcac gaccggcgct tcgccggcga cgcatggcgc accaacctcc catatcgctt

### 4681 cgctgccgcg ttctacctgc tcaatgcgcg cgccttgacc gagctggccg atgccgtcga

### 4741 ggccgatgcc aagacccgcc agcgcatccg cttcgcgatc tcgcaatggg tcgatgcgat

### 4801 gtcgcccgcc aacttccttg ccaccaatcc cgaggcgcag cgcctgctga tcgagtcggg

### 4861 cggcgaatcg ctgcgtgccg gcgtgcgcaa catgatggaa gacctgacac gcggcaagat

### 4921 ctcgcagacc gacgagagcg cgtttgaggt cggccgcaat gtcgcggtga ccgaaggcgc

### 4981 cgtggtcttc gagaacgagt acttccagct gttgcagtac aagccgctga ccgacaaggt

### 5041 gcacgcgcgc ccgctgctga tggtgccgcc gtgcatcaac aagtactaca tcctggacct

### 5101 gcagccggag agctcgctgg tgcgccatgt ggtggagcag ggacatacgg tgtttctggt

### 5161 gtcgtggcgc aatccggacg ccagcatggc cggcagcacc tgggacgact acatcgagca

### 5221 cgcggccatc cgcgccatcg aagtcgcgcg cgacatcagc ggccaggaca agatcaacgt

### 5281 gctcggcttc tgcgtgggcg gcaccattgt ctcgaccgcg ctggcggtgc tggccgcgcg

### 5341 cggcgagcac ccggccgcca gcgtcacgct gctgaccacg ctgctggact ttgccgacac

### 5401 gggcatcctc gacgtctttg tcgacgaggg ccatgtgcag ttgcgcgagg ccacgctggg

### 5461 cggcggcgcc ggcgcgccgt gcgcgctgct gcgcggcctt gagctggcca ataccttctc

### 5521 gttcttgcgc ccgaacgacc tggtgtggaa ctacgtggtc gacaactacc tgaagggcaa

### 5581 cacgccggtg ccgagcgacc tgctgttctg gaacggcgac gccaccaacc tgccggggcc

### 5641 gtggtactgc tggtacctgc gccacaccta cctgcagaac gagctcaagg taccgggcaa

### 5701 gctgaccgtg tgcggcgtgc cggtggacct ggccagcatc gacgtgccga cctatatcta

### 5761 cggctcgcgc gaagaccata tcgtgccgtg gaccgcggcc tatgcctcga ccgcgctgct

### 5821 ggcgaacaag ctgcgcttcg tgctgggtgc gtcgggccat atcgccggtg tgatcaaccc

### 5881 gccggccaag aacaagcgca gccactggac taacgatgcg ctgccggagt cgccgcagca

### 5941 atggctggcc ggcgccatcg agcatcacgg cagctggtgg ccggactgga ccgcatggct

### 6001 ggccgggcag gccggcgcga aacgcgccgc gcccgccaac tatggcaatg cgcgctatcg

### 6061 cgcaatcgaa cccgcgcctg ggcgatacgt caaagccaag gcatga

#### Plasmid Sequences SpyCatch::SpyTag

*pPhaC_Ac_m1::SpyTag003*

### 1 gccaagactc cccgaacgcc gacagacgcc cccgatcgcc ccgcacgcaa ggccggacgt

### 61 ccgacgtccg ccaagccccg cccgcctcgg cagcctaagt cgaaacctgc cgaaacgcgg

### 121 caagccgcct ctgaaaccca tgcagcgccg actcctgaac cgaccgaagc gacacccaat

### 181 caggcccaat tgatcgaaac cctgtcgatg aatctggcca aggcggcaat gatggcccag

### 241 agcgcaatcg ccgaggcggc gctgacccag gccgatcggc cggccgccct ttccgccgat

### 301 cccttcaacg tcgcgccggc catgacgtcg gtcatgacca gcctggccgc ccagcccgac

### 361 aaaatgattc aggcccaggc cgacctgttt ggccgctata tgcagctctg gtcatcgacg

### 421 gcgcgtcagg cggccggcga ggcgcccgat cccgccccgg tcgacaagcg gttcaaggac

### 481 ccggcttggt ccgaaaaccc catgttcgac atgatgcggc gatcctatct gctgacgtcg

### 541 gactggatga acggtctgat cgccggggtc gaggacgtcg atccgacgct gaaacgtcgc

### 601 gcccaattct ttacccgcct gctgaccgac gccttctcgc cgtccaactt tttggcctcc
### 661 aaccccgtcg cgctgaaggc cttggcggag accagcggtg aatcgctggt gaaaggtatg
### 721 cagaacttcg ccgccgactt ggaacgtggc ggcggatcgc tgcggatcag ccaggccgac
### 781 tacggaaagt tcgtcgtcgg cgagaacgtg gccaccgcgc cgggccaggt cgtctggcgc
### 841 gatgagctgt tcgaactgat ccagtacgca ccctcgactg agagtcagca cgacatcccg
### 901 ctgctgatct tcccgccctg gatcaacaaa ttctacatca tggacctgca gccggcgaac
### 961 tcgctgatcc gctggctgtc ggctcagggc ttcacggtct tcgtctgttc ctgggtcaat
### 1021 ccggacaggg ataaggccgg attcggcttc gacgactatc tggacaaggg catctatcgc
### 1081 gcggtcgaga agacgctgga gcaggctggc acgaagcaac tgaacgccgt gggctattgc
### 1141 atcggcggca ccctgcttgg cgctggcctg gcgcacatgg cggccaaggg cgacaagcgc
### 1201 atcgccgccg ctaccttctt cgccgctcaa catgacttcg ccgaggccgg cgacctgctt
### 1261 ctgttcaccg acgaacactg gatcgccgag atcgagcgtc agatggatgc ggcgggcggc
### 1321 gtcctgccgg gcgcggccat ggcggaaacg ttcaacgccc tgcggtccaa tgacctgatc
### 1381 tggtcctttt tcattagcaa ctatctgctg gggaaggatc cgcccgcctt cgacctgttg
### 1441 ttctggaacg ccgatcagac ccgcatgccc aaggcgctgc atctggacta tctgcgtcag
### 1501 atgtatgggg ccaacgccct ggccaagggg cagttcgaga tcggcggcct gacggcggac
### 1561 ctgtcgaagg tcgagatccc gctctatttt caggccagcc gagaggatca catcgcgccg
### 1621 atgaactcgg tctatcgatc cgccaaactg ttcggcggca aggatgtgac cttcaccctg
### 1681 gccgggtcag gtcacatcgc cggcgtcatc aacgcccccg ccgcaaagaa gtatcagcat
### 1741 tggaccaacc ctgccctgcc cgcgaccttg gccgaatggc aagctgacgc cgtcgaacat
### 1801 cccggcagct ggtgggagca ttggtcggcc tggctcggcg cgcgttccgg cgctcaaatc
### 1861 cccgcccgca accccgccaa gggcccgctc aagcccatcg agccggcgcc gggcagctat
### 1921 gtgaaagtga agtcttgaat tatctagagg atccccgggt accgagctcg aattcgcgcg
### 1981 gccgcggcct aggcggcctc ctgtgtgaaa ttgttatccg ctttaattaa aggcatcaaa

### 2041 taaaacgaaa ggctcagtcg aaagactggg cctttcgttt tatctgttgt ttgtcggtga

### 2101 acgctctcct gagtaggaca aatccgccgc cctagacagc tgggcgcgcc ccccctacgg

### 2161 gcttgctctc cgggcttcgc cctgcgcggt cgctgcgctc ccttgccagc ccgtggatat

### 2221 gtggacgatg gccgcgagcg gccaccggct ggctcgcttc gctcggcccg tggacaaccc

### 2281 tgctggacaa gctgatggac aggctgcgcc tgcccacgag cttgaccaca gggattgccc

### 2341 accggctacc cagccttcga ccacataccc accggctcca actgcgcggc ctgcggcctt

### 2401 gccccatcaa tttttttaat ttctctgggg gaaaagcctc cggcctgcgg cctgcgcgct

### 2461 tcgcttgccg gttggacacc aagtggaagg cgggtcaagg ctcgcgcagc gaccgcgcag

### 2521 cggcttggcc ttgacgcgcc tggaacgacc caagcctatg cgagtggggg cagtcgaagg

### 2581 gcgaagcccg cccgcctgcc ccccgagcct cacggcggcg agtgcggggg ttccaagggg

### 2641 gcagcgccac cttgggcaag gccgaaggcc gcgcagtcga tcaacaagcc ccggaggggc

### 2701 cactttttgc cggagggggа gccgcgccga aggcgtgggg gaaccccgca ggggtgccct

### 2761 tctttgggca ccaaagaact agatataggg cgaaatgcga aagacttaaa aatcaacaac

### 2821 ttaaaaaagg ggggtacgca acagctcatt gcggcacccc ccgcaatagc tcattgcgta

### 2881 ggttaaagaa aatctgtaat tgactgccac ttttacgcaa cgcataattg ttgtcgcgct

### 2941 gccgaaaagt tgcagctgat tgcgcatggt gccgcaaccg tgcggcaccc ctaccgcatg

### 3001 gagataagca tggccacgca gtccagagaa atcggcattc aagccaagaa caagcccggt

### 3061 cactgggtgc aaacggaacg caaagcgcat gaggcgtggg ccgggcttat tgcgaggaaa

### 3121 cccacggcgg caatgctgct gcatcacctc gtggcgcaga tgggccacca gaacgccgtg

### 3181 gtggtcagcc agaagacact ttccaagctc atcggacgtt ctttgcggac ggtccaatac

### 3241 gcagtcaagg acttggtggc cgagcgctgg atctccgtcg tgaagctcaa cggccccggc

### 3301 accgtgtcgg cctacgtggt caatgaccgc gtggcgtggg gccagccccg cgaccagttg

### 3361 cgcctgtcgg tgttcagtgc cgccgtggtg gttgatcacg acgaccagga cgaatcgctg

### 3421 ttggggcatg gcgacctgcg ccgcatcccg accctgtatc cgggcgagca gcaactaccg

### 3481 accggccccg gcgaggagcc gcccagccag cccggcattc cgggcatgga accagacctg

### 3541 ccagccttga ccgaaacgga ggaatgggaa cggcgcgggc agcagcgcct gccgatgccc

### 3601 gatgagccgt gttttctgga cgatggcgag ccgttggagc cgccgacacg ggtaacgctg

### 3661 ccgcgccggt agggccggcc tacggccagc ctcgcagagc aggattcccg ttgagcaccg

### 3721 ccaggtgcga ataagggaca gtgaagaagg aacacccgct cgcgggtggg cctacttcac

### 3781 ctatcctgcc cggctgacgc cgttggatac accaaggaaa gtctacacga accctttggc

### 3841 aaaatcctgt atatcgtgcg aaaaaggatg gatataccga aaaaatcgct ataatgaccc

### 3901 cgaagcaggg ttatgcagcg gaaaaggaca acgcgcggac cgcggtccaa ttaattatta

### 3961 gaaaaattca tccagcatca gatgaaattg cagtttgttc atatccggat tatcaatgcc

### 4021 atatttctga aacagacgtt tttgcaggct cgggctaaat tcgcccaggc agttccacag

### 4081 aatggccaga tcctgataac gatccgcaat gcccacacgg cccacatcaa tgcagccaat

### 4141 cagtttgcct tcatcgaaaa tcaggttatc caggctaaaa tcgccgtggg tcaccacgct

### 4201 atccgggcta aacggcagca gtttatgcat ttctttccac acctgttcca ccggccagcc

### 4261 gttacgttca tcatcaaaat cgctcgcatc caccaggccg ttgttcatac ggctctgcgc

### 4321 ctgggccaga cgaaacacac gatcgctgtt aaacgggcag ttgcacaccg gaatgctatg

### 4381 cagacgacgc agaaacacgg ccagcgcatc cacaatgttt tcgccgctat ccggatattc

### 4441 ttccagcacc tgaaacgcgg ttttgcccgg aatcgcggtg gtcagcagcc acgcatcatc

### 4501 cggggtgcga ataaaatgtt taatggtcgg cagcggcata aattcggtca gccagttcag

### 4561 acgcaccatt tcatcggtca catcgttcgc cacgctgcct ttgccatgtt tcagaaacag

### 4621 ttccggcgca tccggtttgc catacagacg ataaatggtc gcgccgctct gacccacgtt

### 4681 atcacgcgcc catttatagc catacagatc cgcatccatg ttgctgttca gacgcggacg

### 4741 gctacagctc gtttcacgct gaatatggct cataacaccc cttgtattac tgtttatgta

### 4801 agcagacagt tttattgttc atgatgatat atttttatct tgtgcaatgt aacatcagag

### 4861 attttgagac acaaatttaa atcgtaatta ttggggaccc ctggattctc accaataaaa

### 4921 aacgcccggc ggcaaccgag cgttctgaac aaatccagat ggagttctga ggtcattact

### 4981 ggatctatca acaggagtcc aagactagtc gccagggttt tcccagtcac gacgcggccg

### 5041 caagcttgca tgcctgcagg ttatgacaac ttgacggcta catcattcac tttttcttca

### 5101 caaccggcac ggaactcgct cgggctggcc ccggtgcatt ttttaaatac ccgcgagaaa

### 5161 tagagttgat cgtcaaaacc aacattgcga ccgacggtgg cgataggcat ccgggtggtg

### 5221 ctcaaaagca gcttcgcctg gctgatacgt tggtcctcgc gccagcttaa gacgctaatc

### 5281 cctaactgct ggcggaaaag atgtgacaga cgcgacggcg acaagcaaac atgctgtgcg

### 5341 acgctggcga tatcaaaatt gctgtctgcc aggtgatcgc tgatgtactg acaagcctcg

### 5401 cgtacccgat tatccatcgg tggatggagc gactcgttaa tcgcttccat gcgccgcagt

### 5461 aacaattgct caagcagatt tatcgccagc agctccgaat agcgcccttc cccttgcccg

### 5521 gcgttaatga tttgcccaaa caggtcgctg aaatgcggct ggtgcgcttc atccgggcga

### 5581 aagaaccccg tattggcaaa tattgacggc cagttaagcc attcatgcca gtaggcgcgc

### 5641 ggacgaaagt aaacccactg gtgataccat tcgcgagcct ccggatgacg accgtagtga

### 5701 tgaatctctc ctggcgggaa cagcaaaata tcacccggtc ggcaaacaaa ttctcgtccc

### 5761 tgatttttca ccaccccctg accgcgaatg gtgagattga gaatataacc tttcattccc

### 5821 agcggtcggt cgataaaaaa atcgagataa ccgttggcct caatcggcgt taaacccgcc

### 5881 accagatggg cattaaacga gtatcccggc agcaggggat cattttgcgc ttcagccata

### 5941 cttttcatac tcccgccatt cagagaagaa accaattgtc catattgcat cagacattgc

### 6001 cgtcactgcg tcttttactg gctcttctcg ctaaccaaac cggtaacccc gcttattaaa

### 6061 agcattctgt aacaaagcgg gaccaaagcc atgacaaaaa cgcgtaacaa aagtgtctat

### 6121 aatcacggca gaaaagtcca cattgattat ttgcacggcg tcacactttg ctatgccata

### 6181 gcatttttat ccataagatt agcggatcct acctgacgct ttttatcgca actctctact

### 6241 gtttctccat gggccccccc aaaggaggac aaccatggta accaccttat caggtttatc

### 6301 aggtgagcaa ggtccgtccg gtgatatgac aactgaagaa gatagtgcta cccatattaa

### 6361 attctcaaaa cgtgatgagg acggccgtga gttagctggt gcaactatgg agttgcgtga

### 6421 ttcatctggt aaaactatta gtacatggat ttcagatgga catgtgaagg atttctacct

### 6481 gtatccagga aaatatacat ttgtcgaaac cgcagcacca gacggttatg aggtagcaac

### 6541 tccaattgaa tttacagtta atgaggacgg tcaggttact gtagatggtg aagcaactga

### 6601 aggtgacgct catactggat ccagtggtag c
